# Totum-448, a polyphenol-rich plant extract, decreases hepatic steatosis and inflammation in diet-induced MASLD mice

**DOI:** 10.1101/2025.03.24.644956

**Authors:** Joost M. Lambooij, Vivien Chavanelle, Marie Vallier, Hendrik J.P. van der Zande, Yolanda F. Otero, Frank Otto, Robbie Schuurman, Florian Le Joubioux, Thierry Maugard, Martin Giera, Sébastien L. Peltier, Arnaud Zaldumbide, Pascal Sirvent, Bruno Guigas

## Abstract

The increasing prevalence of obesity-driven metabolic dysfunction-associated steatotic liver disease (MASLD) urges the development of new therapeutic strategies. Totum-448 is a unique patented combination of polyphenol-rich plant extracts designed to reduce hepatic steatosis, a risk factor for steatohepatitis and type 2 diabetes. In this study, we investigated the effects of Totum-448 on metabolic homeostasis and steatohepatitis in diet-induced MASLD mice. For this purpose, male C57Bl6/J mice were fed a high-fat diet in combination with sucrose-containing drinking water for 12 weeks, followed by diet supplementation with or without Totum-448 for 4 weeks. Body weight/composition, caloric intake, plasma parameters and whole-body glucose tolerance were measured throughout the study and fecal microbiota composition was determined by 16S sequencing. Hepatic steatosis, transcriptomic/lipidomic profiles and immune cell composition were assessed by histological/biochemical assays, RNA sequencing, MS-based lipidomics, and spectral flow cytometry. We found that Totum-448 significantly lowered hyperinsulinemia and systemic glucose intolerance in MASLD mice without affecting body weight, fat mass, calorie intake, feces production or fecal microbiota composition. Furthermore, a decrease in liver MASLD activity score and macrovesicular steatosis, hepatic triglycerides and cholesterol contents, and plasma alanine aminotransferase levels were observed. Totum-448 also reduced the liver inflammatory and pro-fibrotic transcriptomic signatures and decreased both MASLD-induced loss in tissue-resident Kupffer cells and recruitment of monocyte-derived pro-inflammatory macrophages. Altogether, Totum-448 reduces hepatic steatosis and inflammation in insulin-resistant, steatotic, obese mice, a dual effect that likely contributes to improved whole-body metabolic homeostasis. Supplementation with Totum-448 may therefore constitute a promising novel nutritional approach for MASLD patients.

## Introduction

The development of metabolic dysfunction-associated steatotic liver disease (MASLD), formerly known as non-alcoholic fatty liver disease (NAFLD)^1^, and its progressive more aggressive form, metabolic dysfunction-associated steatohepatitis (MASH), are closely intertwined with the current worldwide epidemic of obesity and type 2 diabetes^2,3^. Recent epidemiological studies have indeed highlighted the alarming rise of MASLD in both developing and developed countries, with a high prevalence rate (>50%) in overweight and obese adults that constitutes one of the health challenges of the 21^st^ century^4^.

MASLD has a complex pathophysiology, characterized by hepatic lipid accumulation, insulin resistance, lipotoxicity, inflammation, and progressive fibrogenesis^1^, with an estimated annual cost exceeding 200 billion euros in the United States and Europe alone^5^. The disease spectrum is broad, ranging from isolated steatosis to MASH, fibrosis, and cirrhosis, ultimately increasing the risk of hepatocellular carcinoma^6–8^. Effective therapeutic treatments remain scarce as only Resmetirom, a modestly effective drug for fibrotic MASH, was recently approved by the FDA^9^.

Metaflammation, which refers to a state of chronic, low-grade inflammation arising from obesity-associated immunometabolic dysregulations in various organs, plays a pivotal role in MASLD initiation and progression^10^. Excessive hepatic influx of lipids and accumulation of triglycerides in form of lipid droplets in hepatocytes triggers a cascade of inflammatory responses in the liver mediated by various immune cell types, notably those from the innate myeloid compartment^11,12^. In this context, macrophages are believed to play a central role^13–16^. During homeostasis, hepatic macrophages predominantly consist of self-replenishing, embryonically-derived tissue-resident Kupffer cells (resKCs)^17,18^. However, in response to obesity-associated lipotoxic stress and local inflammatory cues in their microenvironment, resKCs undergo cell death, leading to an influx of circulating bone-marrow-derived monocytes for replenishing the empty niche. These cells further differentiate into monocyte-derived macrophages (moMACS) and ultimately resKCs that display almost identical features than KCs of embryonic origin^19,20^. Collectively, these obesogenic-driven changes in the hepatic immunological landscape contribute to the chronic inflammatory milieu, leading to liver injury, fibrosis, and ultimately, the development of MASH and its complications^21–23^. Moreover, metaflammation extends beyond the liver, promoting systemic metabolic dysfunction and exacerbating both central and peripheral insulin resistance, dyslipidemia, and cardiovascular risk^24^.

The multifactorial nature of MASLD highlights the need for comprehensive strategies involving conventional lifestyle interventions and innovative preventive and/or therapeutic approaches to halt disease progression and reduce associated morbidity and mortality^25^. Recently, functional foods and nutraceuticals containing various bioactive compounds have received considerable attention due to their potential therapeutic benefits in the context of MASLD/MASH^26,27^. Indeed, their accessibility and relatively low risk of adverse effects make them attractive adjuncts to lifestyle modifications and/or pharmacological treatments. For instance, bioactive compounds such as omega-3 fatty acids, polyphenols, flavonoids, and vitamins, as well as probiotics, have been shown to mitigate lipid accumulation, inflammation, oxidative stress and insulin resistance in the liver, potentially contributing to prevent MASLD/MASH progression and/or promoting disease regression^28^.

Totum-448 is a novel, polyphenol-rich plant-based active principle composed of a mixture of plant extracts designed to reduce obesity-induced hepatic steatosis, a risk factor for progression towards type 2 diabetes and MASH. In the present study, we aimed to investigate the effects of Totum-448 on hepatic steatosis, liver inflammation and metabolic homeostasis in a dietary mouse model of MASLD.

## Materials and Methods

### Totum-448

Totum-448 is a patented blend of 5 plant extracts and choline designed to act in combination to target the risk factors of developing MASLD. The mixture contains extracts from olive leaf (*Olea europaea*), bilberry (*Vaccinium myrtillus*), artichoke leaf (*Cynara scolymus*), chrysanthellum (*Chrysanthellum indicum* subsp*. afroamericanum B.L. Turner*), black pepper (*Piper nigrum*) and choline. **Table S1** shows the chemical characterization of Totum-448.

### Animals and diet

All experiments were performed in accordance with the Guide for the Care and Use of Laboratory Animals of the Institute for Laboratory Animal Research and were approved by the Dutch ethical committee on animal experiments (Centrale Commissie Dierproeven; AVD1060020174364). An *a priori* power calculation was done. Ten-week-old C57BL/6JOlaHsd male mice were purchased from Envigo (Horst, The Netherlands) and housed in a temperature-controlled room with a 12-hour light-dark cycle and *ad libitum* access to food and drink. Mice were fed a low-fat diet (LFD, 10% energy derived from fat, D12450H, Research Diets, New Brunswick, NJ, USA) or high fat diet (HFD, 45% energy derived from fat, D12451, Research Diets, New Brunswick, NJ, USA) supplemented with sucrose in the drinking water (10% w/v, HFD/S) for 12 weeks. The experimental groups were randomized after removal of HFD/S low responders (∼5%; body weight gain <6 g), after which HFD was supplemented either with Totum-448 (Valbiotis SA, Perigny, France) or not for an additional 4 weeks. The experimenters were not blinded to the diet supplementation on the metabolic test days, however, most of the subsequent analyses were performed in blind conditions.

### Body composition, energy intake and feces production

Body weight was frequently measured during the 4 weeks of supplementation using a conventional weighing scale. Body composition was measured by MRI (Echo Medical Systems, Houston, TX, USA) in conscious unrestrained mice. At sacrifice, visceral white adipose tissue (epidydimal; eWAT), supraclavicular brown adipose tissue (BAT), heart and liver were weighed and collected for further processing. The intestines were collected and measured (total and colon separately) and the weight of the cecum was determined using a precision scale. Food and sucrose intake were frequently assessed throughout the study by weighing food pellets and measuring liquid volume in drinking bottle for every cage (2-3 mice per cage). At week 4, feces produced over 24h were carefully collected in cage bedding and weighed.

### Glucose tolerance test

Whole-body intraperitoneal (i.p.) glucose tolerance (ipGTT) test was performed at week 4 of Totum-448 supplementation, as previously reported^29^. In short, a bolus of glucose (2g D-glucose/kg body weight; Sigma-Aldrich, St. Louis, MO, USA) was administered i.p. in 6h-fasted mice and blood glucose was measured at t=0, 20, 40, 60, and 90 min post glucose injection using a Glucometer (Accu-Check; Roche Diagnostics, Basel, Switzerland).

### Plasma analysis

Blood samples were collected from the tail vein of 4h-fasted mice using paraoxon-coated glass capillaries. Plasma insulin was determined using a commercially available ELISA kit (Chrystal Chem, Elk Grove Village, IL, USA) according to the manufacturer’s instructions. The homeostatic model assessment of insulin resistance (HOMA-IR) adjusted for mice^30^ was calculated as followed ([glucose (mg/dl)*0.055]*[insulin (ng/ml)*172.1])/3875. Plasma alanine aminotransferase (ALAT) was measured using a Reflotron® kit (Roche diagnostics, Basel, Switzerland).

### Fecal microbiota analyses

DNA was extracted from fecal samples using the FastDNA^TM^ Spin Kit for Feces and a FastPrep-24 5G (MP Biomedicals, Santa Ana, CA, USA) following the manufacturer’s instructions. Microbial 16S library preparation was performed at the PGTB (Plateforme Génome Transcriptome de Bordeaux, Bordeaux, France) by amplification and sequencing of the V3-V4 region of the 16S rRNA gene on an Illumina MiSeq using the 2×250bp Illumina v2 kit (Illumina, San Diego, CA, USA). Data processing and statistical analyses are described in details in the supplementary methods section.

### Hepatic lipid composition

Liver triglycerides (TG), total cholesterol (TC) and phospholipid (PL) contents were measured using commercial kits (Instruchemie, Delfzijl, The Netherlands) #2913, #10015 and #3009, respectively) and expressed as nmol per mg of total protein content using the Bradford protein assay kit (Sigma-Aldrich, St. Louis, MO, USA), as previously reported^31,32^. For lipidomics, lipids were extracted from 10 mg of liver by the methyl-tert-butylether method and analyzed using the Lipidyzer™, a direct infusion-tandem mass spectrometry-based platform (Sciex, Redwood City, CA, USA), as previously described^32^. Lipid concentrations were expressed as pmol/mg of liver.

### Histological analysis

Pieces of liver (∼30 mg) were fixed in 4% formaldehyde (Sigma-Aldrich, St. Louis, MO, USA), paraffin-embedded, sectioned at 4 μm and stained with Hematoxilin and Eosin (H&E). After scanning, 5 fields at 40x or 20x magnification were used for the determination of lipid droplet (LD) size distribution and mean area, and MASLD activity score (NAS), respectively, as previously reported^33^.

### Isolation of blood and liver leukocytes for flow cytometry

At sacrifice, blood was collected retro-orbitally in heparin-coated tubes for leukocyte isolation, as described previously^29^ and briefly described below. For liver samples, the organs were collected after a 1 min post-sacrifice transcardial perfusion with PBS and further digested for isolation of leukocytes, as previously reported^34^ and briefly described below.

#### Blood

Samples were diluted 1:1 in PBS (Fresenius Kabi, Bad Homburg, Germany, with calcium and magnesium) and erythrocytes were lysed for 15 min at room temperature using an erythrocyte lysis/fixation solution (BD Biosciences, Franklin Lakes, NJ, USA). Leukocytes were then centrifuged at 500 x g for 5 min at 4°C and then subsequently washed three times in PBS. After washing, cell pellets were resuspended in PBS supplemented with 1% heat inactivated fetal calf serum (hiFSC; Serana, Pessin, Germany) and 2.5 mM ethylenediaminetetraacetic acid (EDTA; Sigma-Aldrich, St. Louis, MO, USA), counted using a hemocytometer and 1*10^6^ cells per sample were further processed for flow cytometry.

#### Liver

Livers were first collected in 10 mL ice-cold RPMI 1640 + Glutamax (Thermo Fisher Scientific, Waltham, MA, USA), minced and digested for 25 min at 37°C under agitation (200 RPM) in 5 mL RPMI 1640 + Glutamax supplemented with 1 mg/ml Collagenase Type IV from *Clostridium histolyticum* (Sigma-Aldrich, St. Louis, MO, USA, 125 CDU/ml), 1 mg/ml Dispase II (Sigma-Aldrich, St. Louis, MO, USA, 1.4 U/ml), 1 mg/ml Collagenase D from *C. histolyticum* (Roche, Basel, Switzerland, 250 Mandl U/ml) and 2000 U/mL DNase I (Sigma-Aldrich, St. Louis, MO, USA). After digestion, samples were filtered (100 µM cell strainer; Corning, NY, USA) and pelleted at 300 x g for 10 min at 4°C after which the pellets were washed twice with 40 mL PBS/hiFSC/EDTA. After washing, the pellets were treated with 3 mL erythrocyte lysis buffer consisting of 0.15 M NH4Cl (Merck, Rahway, NJ, USA), 1 mM KHCO3 (Merck, Rahway, NJ, USA) and 0.1 mM EDTA in ddH2O for 2 min at room temperature. Next, total leukocytes were isolated by MACS-sorting using CD45 positive selection MicroBeads and LS columns (Mitenyi Biotec, Bergisch Gladback, USA) according to the manufacturer’s instructions. Post-isolation, total leukocytes were counted using a hemacytometer and 1*10^6^ cells per sample were further processed for flow cytometry.

### Flow cytometry

#### Blood

Isolated blood leukocytes were washed with PBS/hiFSC/EDTA, pelleted at 500 x g for 5 min at 4°C and incubated with a cocktail of antibodies directed against CD3, CD4, CD8, CD11b, CD19, CD45, Ly6C, NK1.1 and Siglec-F (see **Table S3** for details) in PBS/hiFSC/EDTA supplemented with Brilliant Stain Buffer Plus (BD Biosciences, Franklin Lakes, NJ, USA) and True Stain Monocyte Blocker (Biolegend, San Diego, CA, USA) for 30 min at room temperature. After washing, the cells were resuspended in PBS/hiFSC/EDTA and acquired on a 3-laser Cytek Aurora (Cytek Biosciences, Fremont, CA, USA).

#### Liver

Isolated liver leukocytes were pelleted at 500 x g for 5 min at 4°C and subsequently incubated with Zombie-NIR viability dye in PBS supplemented with True Stain Monocyte Blocker for 20 min at room temperature. After washing with PBS and pelleting as described above, the cells were fixed using a 2% paraformaldehyde solution (PFA; Sigma-Aldrich, St. Louis, MO, USA) in PBS for 10 min at room temperature. Post-fixation, the cells were washed with PBS/hiFSC/EDTA and incubated with a cocktail of antibodies directed against CD3, CD11b, CD11c, CD19, CD45, CD64, CD90.2, CLEC2, F4/80, Ly6C, Ly6G, NK1.1, Siglec-F, TIM4 and TREM2 (see **Table S3** for details) in PBS/hiFCS/EDTA supplemented with Brilliant Stain Buffer Plus and True Stain Monocyte Blocker for 30 min at 4°C. After washing, the cells were resuspended in PBS/FSC/EDTA and acquired on a 5-laser Cytek Aurora (Cytek Biosciences, Fremont, CA, USA).

SpectroFlo v3.0 (Cytek Biosciences, Fremont, CA, USA) was used for spectral unmixing and FlowJo^TM^ v10.8 was used to gate the flow cytometry data for all samples. Representative gating strategies used to gate the blood and liver cells can be found in **Fig. S3a** and **Fig. S4a**, respectively.

### RNA isolation and RNA sequencing analyses

RNA was extracted from snap-frozen liver samples (∼10-20 mg) using an RNA purification kit (NucleoSpin RNA Midi, Macherey-Nagel, Düren, Germany) followed by an on-filter DNAse treatment, according to the instructions provided by the manufacturer. RNA concentration and purity were measured using NanoDrop 2000 (Thermo Fisher Scientific, Waltham, MA, USA). RNA integrity was assessed using the RNA Nano 6000 Assay Kit of the Agilent Bioanalyzer 2100 system (Agilent Technologies, Santa Clara, CA, USA). Further RNA sample preparation, sequencing and data pre-processing were done at Biomarker Technologies (BMK GmbH Münster, Germany), as described in the supplementary method section.

### RNA isolation and RT-qPCR

RNA was extracted from snap-frozen liver (∼10-20 mg) using TriPure RNA Isolation reagent (Roche Diagnostics, Basel, Switzerland). Total RNA (1-2 μg) was reverse transcribed using the M-MLV Reverse Transcriptase kit (Thermo Fisher Scientific, Waltham, MA, USA). Real-time qPCR runs were performed on a CFX96 Real-time C1000 thermal cycler (Biorad, Hercules, CA, USA) using the GoTaq qPCR Master Mix kit (Promega, Madison, WI, USA). Gene expression was normalized using housekeeping gene *RplP0* and expressed as fold change compared to LFD-fed mice. Primer sequences can be found in **Table S2**.

### Statistical analysis

All data are presented as mean ± standard error of the mean (SEM). Statistical analysis was performed using GraphPad Prism 8.0 (GraphPad Software, La Jolla, CA, USA) with unpaired t-test, one-way or two-way analysis of variance (ANOVA) followed by Fisher’s post-hoc test. Differences between groups were considered statistically significant at p < 0.05. Outliers were identified according to the two-standard deviation method (GraphPad Software, La Jolla, CA, USA).

## Results

### Totum-448 improves whole-body metabolic homeostasis in MASLD mice independently of body weight changes

To select the optimal Totum-448 concentration to be administered, a pilot study was performed in MASLD mice (**Fig. S1**). For this purpose, C57BL/6 male mice were first fed a high-fat diet supplemented with sucrose in the drinking water (HFD/S) for 12 weeks, followed by HFD supplementation with or without Totum-448 at various concentrations (1.5, 2 and 2.5% w/w) for 4 additional weeks (**Fig. S1a**). We observed a substantial time- and dose-dependent decrease in body weight at the two highest Totum-448 concentrations (−3.8% and −9.8% at 2 and 2.5% Totum-448, respectively (**Fig. S1b-c**)). These effects were likely due to a dose-dependent decrease in food intake (**Fig. S1d**) even though concomitant increases in liquid energy intake were evidenced (**Fig. S1e**). Altogether, no significant impact on total energy intake (**Fig. S1f**) and feces production (**Fig. S1g**) were observed. To further study the effects of Totum-448 on metabolic homeostasis in insulin resistant obese MASLD mice, the concentration that did not affect body weight and food intake was selected, *i.e.* 1.5% w/w, and administered to HFD/S-fed mice using the same experimental settings as described above (**Fig. 1a**). In line with the pilot study, we did not observed any effect on body weight (**Fig. 1b**) and composition (**Fig. 1c-d**) in obese mice after a 4-week supplementation with 1.5% Totum-448. As expected, HFD/S feeding increased fasting plasma glucose and insulin levels (**Fig. 1e-f**), and HOMA-IR (**Fig. 1g**) when compared to low-fat diet (LFD)-fed mice. Furthermore, HFD/S impaired whole-body glucose homeostasis, as assessed by intraperitoneal glucose tolerance test (**Fig. 1h**). Although no effect was observed on fasting plasma glucose levels, Totum-448 supplementation in HFD/S-fed mice significantly reduced insulin levels, and, consequently, the calculated HOMA-IR (**Fig. 1e-g**). Congruent with HOMA-IR data, Totum-448 improved whole-body glucose homeostasis in obese mice (**Fig. 1h**), without affecting glucose-induced insulin levels (*data not shown*). Of note, these weight change-independent effects of Totum-448 on insulin and HOMA-IR were already observed after 2 weeks of supplementation (**Fig. S2**). In addition, although the HFD/S-induced increase in total blood leukocyte counts was unchanged, the circulating levels of monocytes was significantly reduced by Totum-448, while other myeloid (neutrophils, eosinophils) and lymphoid (B, NK, CD4^+^ and CD8^+^ T) subsets were not affected (**Fig. S3**).

**Figure 1.**
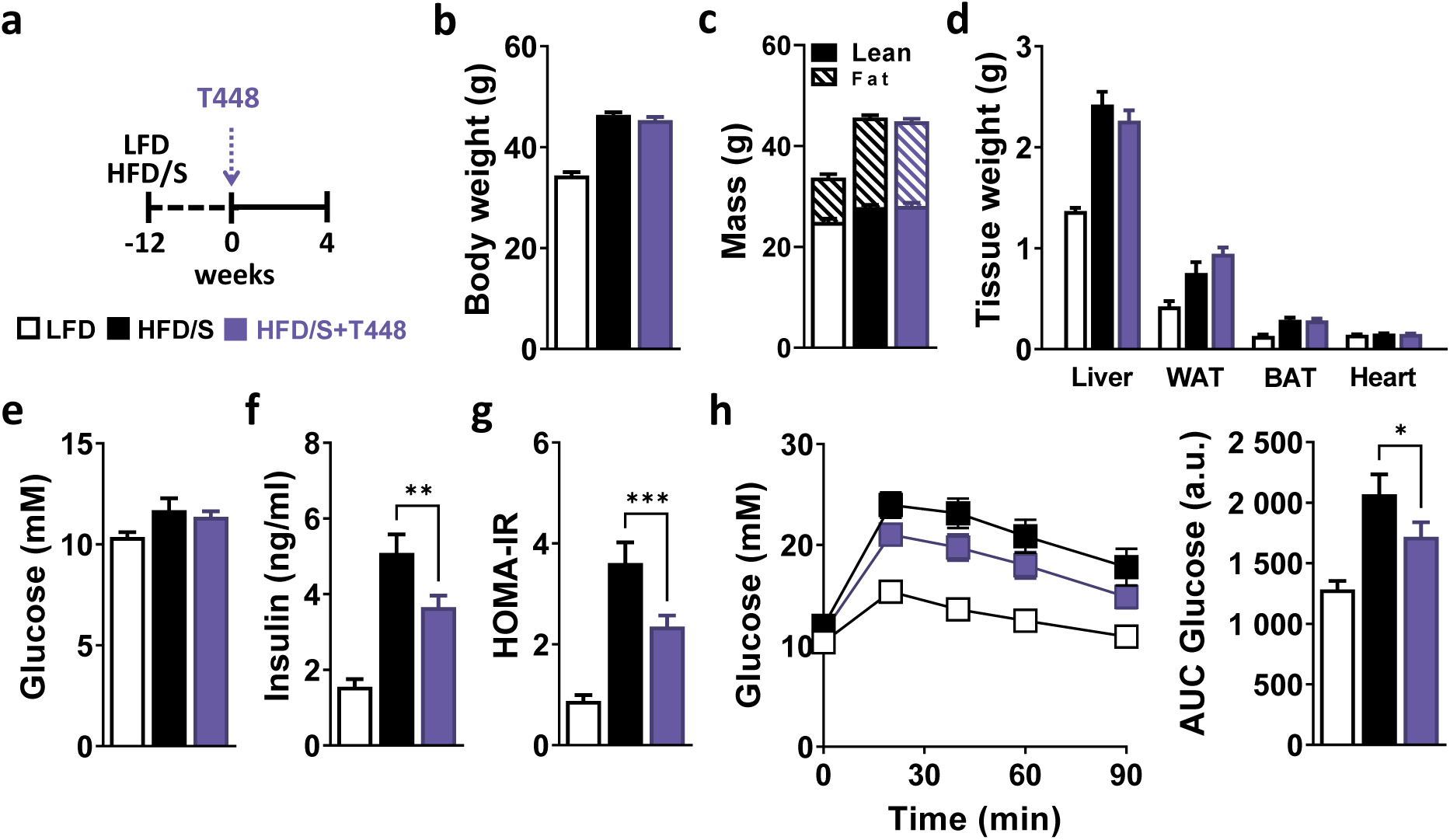
T448 improves HFD-induced insulin resistance without affecting body weight and body composition. 10 week-old C57BL/6JOlaHsd male mice were fed either a low-fat diet (LFD, open squares/bars) or high-fat diet (HFD) supplemented with sucrose in the drinking water (10% w/v, HFD/S) for a period of 12 weeks after which the HFD was either supplemented with Totum-448 (T448, 1.5% g/g; purple squares/bars) or left without supplementation (control; black squares/bars) for 4 additional weeks (**a**). At week 4 of treatment, body weight (**b**) and body composition (**c**) were determined. Post-sacrifice, the weight of the liver, WAT, BAT and heart were determined (**d**). The fasting glucose and insulin levels (**e-f**) were determined at week 4 and used to calculate HOMA-IR (**g**). An intraperitoneal (i.p.) glucose tolerance test (GTT) was performed at week 4 in 6-hour fasted mice. Blood glucose levels were measured at baseline and 20, 40, 60 and 90 min post-injection, and the AUC was calculated (**h**). Results are expressed as mean ± SEM. * p ≤ 0.05, ** p ≤ 0.01, *** p ≤ 0.001. n=10-12 mice per group from 2 independent experiments.

### Totum-448 marginally impacts fecal microbiome composition

Given that obesity-induced changes in gut microbiota is associated with metabolic dysfunctions^35^, we next assessed the impact of Totum-448 on fecal microbiome composition by performing 16S ribosomal RNA sequencing on feces collected during the last week of the study. Importantly, Totum-448 supplementation had no effect on the length of total intestine and colon nor the weight of the cecum content at sacrifice (**Fig. 2a-c**). At the phylum level, the Shannon index was decreased in the HFD/S groups when compared to the LFD group, indicating a reduction in bacterial species diversity, but no significant differences were observed in response to Totum-448 supplementation (**Fig. 2d**). Principal component analysis (PCA) of the relative abundance of intestinal microbial communities using the Bray-Curtis dissimilarity index also confirmed that microbial composition only differed significantly between the LFD and HFD/S groups (notably *Actinobacteria* and *Tenericutes*) but not in response to Totum-448 supplementation (**Fig. 2e-f, Table S4**). At the genus level, the Shannon index was increased in both HFD/S and HFD/S+Totum-448 groups (**Fig. 2g**). The PCA analysis confirmed that the HFD/S strongly altered microbial composition when compared to LFD while Totum-448 supplementation had no significant effect (**Fig. 2h**). Similar PCA results were obtained using Jaccard distance (**Fig. 2i**), which is more sensitive to rare taxa by only taking into account the presence or absence of a dedicated taxon, independent of its abundance. Altogether, only few bacterial taxa from the *Firmicutes* phylum were specifically affected by Totum-448 (**Fig. 2j, Table S5**), indicating a marginal impact on fecal microbiota composition.

**Figure 2.**
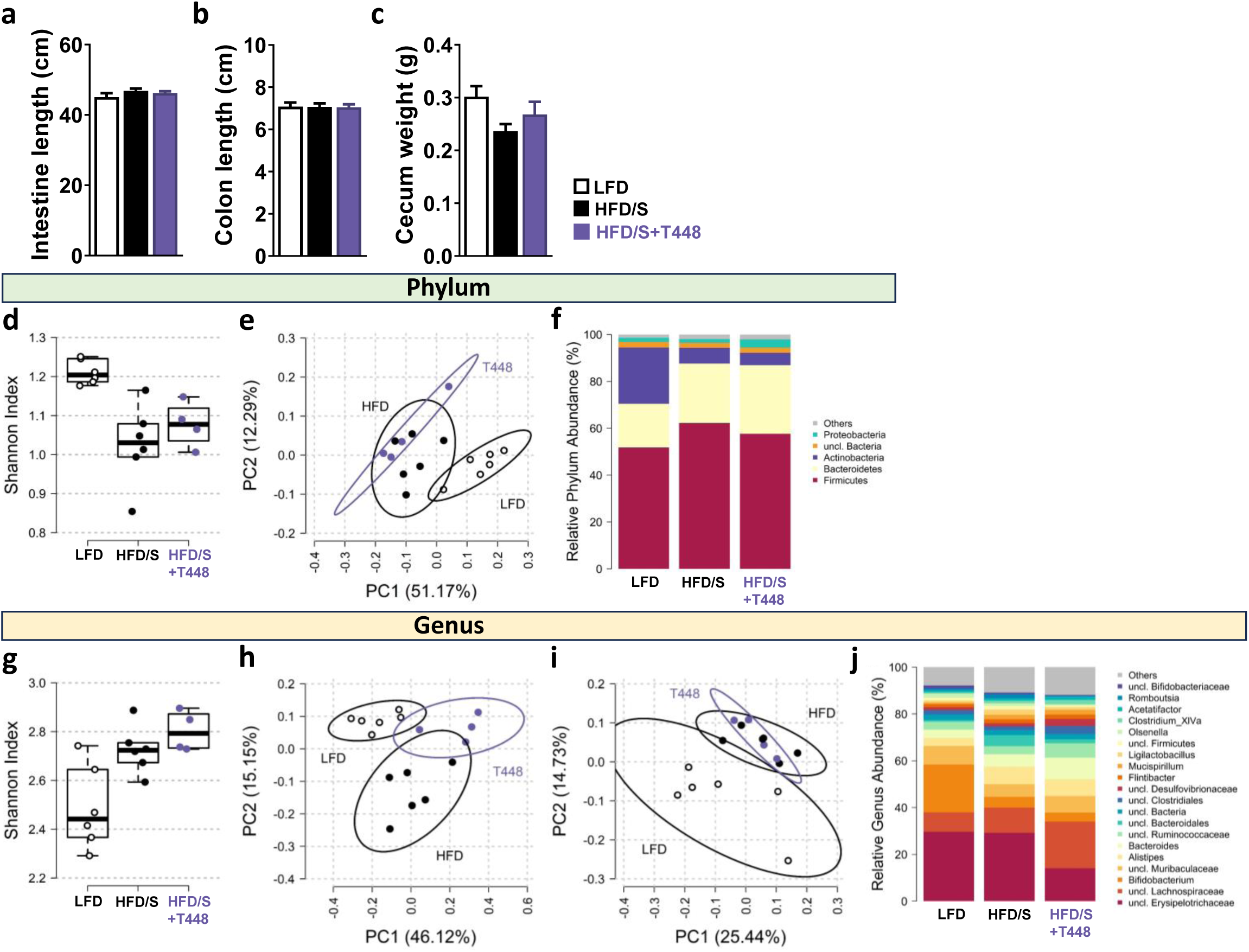
Totum-448 does not have significant impact on intestine length and fecal microbiome composition. LFD- and HFD/S-fed mice were treated as described in Fig. 1. The intestine and colon lengths and the cecum weight were measured post-sacrifice (**a-c**). Fecal microbiome alpha and beta diversity: species richness and evenness were assessed at phylum (**d**) and genus (**g**) level by Shannon index. Principal coordinate analysis of microbial composition using Bray-Curtis (**e,h**) and Jaccard (**i**) distances at phylum (**e**) and genus (**h,i**) levels. Average relative microbiome composition of fecal samples at phylum (**f**) and genus (**j**) levels. For visual clarity, only the most abundant 5 phyla and 20 genera are presented individually, the rest being summed up into “Others”. Results are expressed as mean ± SEM. n=10-12 mice per group from 2 independent experiments.

### Totum-448 reduces hepatic steatosis and alters liver lipid composition

Ectopic lipid accumulation, especially in the liver, triggers immunometabolic dysfunctions contributing to insulin resistance and impaired nutrient homeostasis^7^. Therefore, we next investigated the impact of Totum-448 supplementation on hepatic steatosis in MASLD mice. Remarkably, Totum-448 almost completely reverted HFD/S-induced hepatic steatosis in MASLD mice, as assessed by H&E staining (**Fig. 3a**). This effect was mostly resulting from a reduction in macrovascular steatosis (**Fig. 3b-c**) and associated with a significant reduction of steatosis, inflammation, hepatocellular ballooning and MASLD activity scores (**Fig. 3d**). These findings were further supported by a potent decrease in both liver triglycerides (TG) and total cholesterol contents (−28% and −30% respectively; p<0.05; **Fig. 3e**). Quantitative lipidomics further confirmed that Totum-448 significantly affected the hepatic lipid composition by reducing the liver content of a large numbers of TGs, diglycerides (DGs) and free fatty acid (FFA) species in MASLD mice (**Fig. 3f-g**).

**Figure 3.**
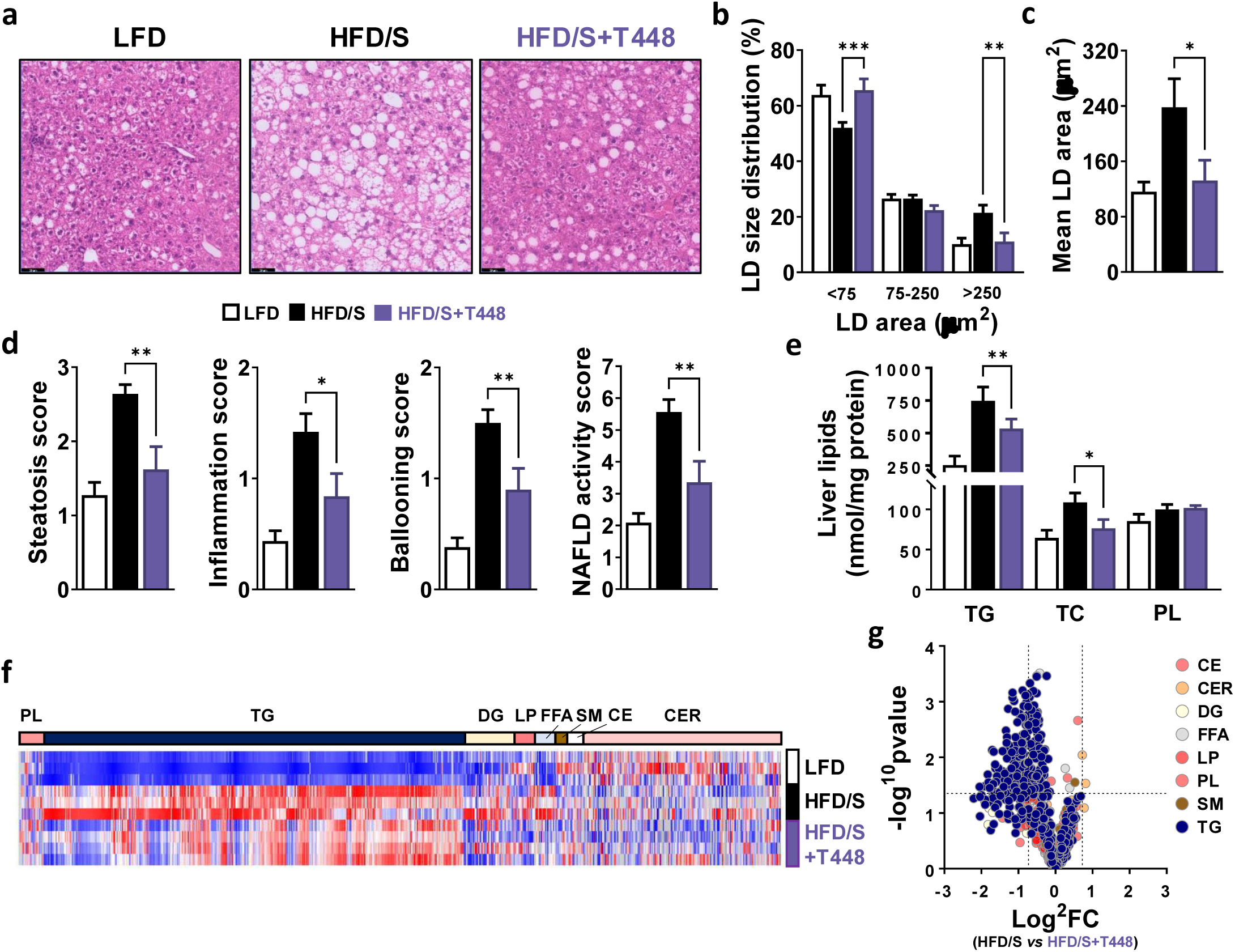
Totum-448 reduces hepatic steatosis. LFD- and HFD/S-fed mice were treated as described in Fig.1. PFA-fixed, paraffin-embedded liver section were stained with Hematoxilin and Eosin (H&E, **a**) followed by computer-assisted determination of hepatic lipid droplet (LD) size distribution (**b**) and mean LD area (**c**). H&E-stained slides were also used to assess the hepatic steatosis, lobular inflammation, hepatocellular ballooning scores and overall MAFLD activity score (NAS) (**d**). Hepatic triglyceride (TG), total cholesterol (TC) and phospholipid (PL) contents (**e**) were determined post-sacrifice. The hepatic lipid composition was determined by targeted lipidomics using the Lipidyzer platform. The heatmap shows the relative abundance of the individual lipid species per class in each group (**f**). The relative increase and decrease of various lipid species per class in livers from HFD/S+T448 compared to HFD/S-fed mice are displayed on the volcano plot (**g**). CE, Cholesterylester; CER, ceramides; DG, Diglycerides; FFA, Free-fatty acids; LP, Lipoprotein; PL, Phospholipids; SM, Sphingomyelin; TG, Triglycerides. Results are expressed as mean ± SEM. * p ≤ 0.05, ** p ≤ 0.01, *** p ≤ 0.001. n=10-12 mice per group from 2 independent experiments for **a-e** and n=3-4 mice per group for lipidomics.

### Totum-448 lowers inflammatory and pro-fibrotic transcriptomic signatures in the liver

To gain mechanistic insights into the beneficial metabolic effects of Totum-448, bulk RNA sequencing was performed in the livers from MASLD mice. Differential gene expression analysis showed that Totum-448 induced a significant up- and downregulation of 47 and 345 unique transcripts, respectively, in HFD/S-fed mice (**Fig. 4a-b**). Gene ontology and gene set enrichment analyses indicated an enrichment of downregulated genes involved in innate immune response, myeloid cell and platelet activation, pro-inflammatory cytokine production and extracellular matrix organization (**Fig. 4c-d**). A large number of genes encoding proteins involved in liver inflammation (*e.g. Lcn2*) and hepatic stellate cell pro-fibrotic activation (*e.g. Acta2*, *Timp1*, *Col1a1* and *Mmp12*) were found to be among the most significantly downregulated by Totum-448. These findings were confirmed by targeted qPCR (**Fig. 4e**). In line with improvements in hepatic MASLD/MASH features, a decrease in circulating alanine aminotransferase (ALT) was observed in response to Totum-448 supplementation, indicating a reduction in hepatocyte injury and liver damage (**Fig. 4f**).

**Figure 4.**
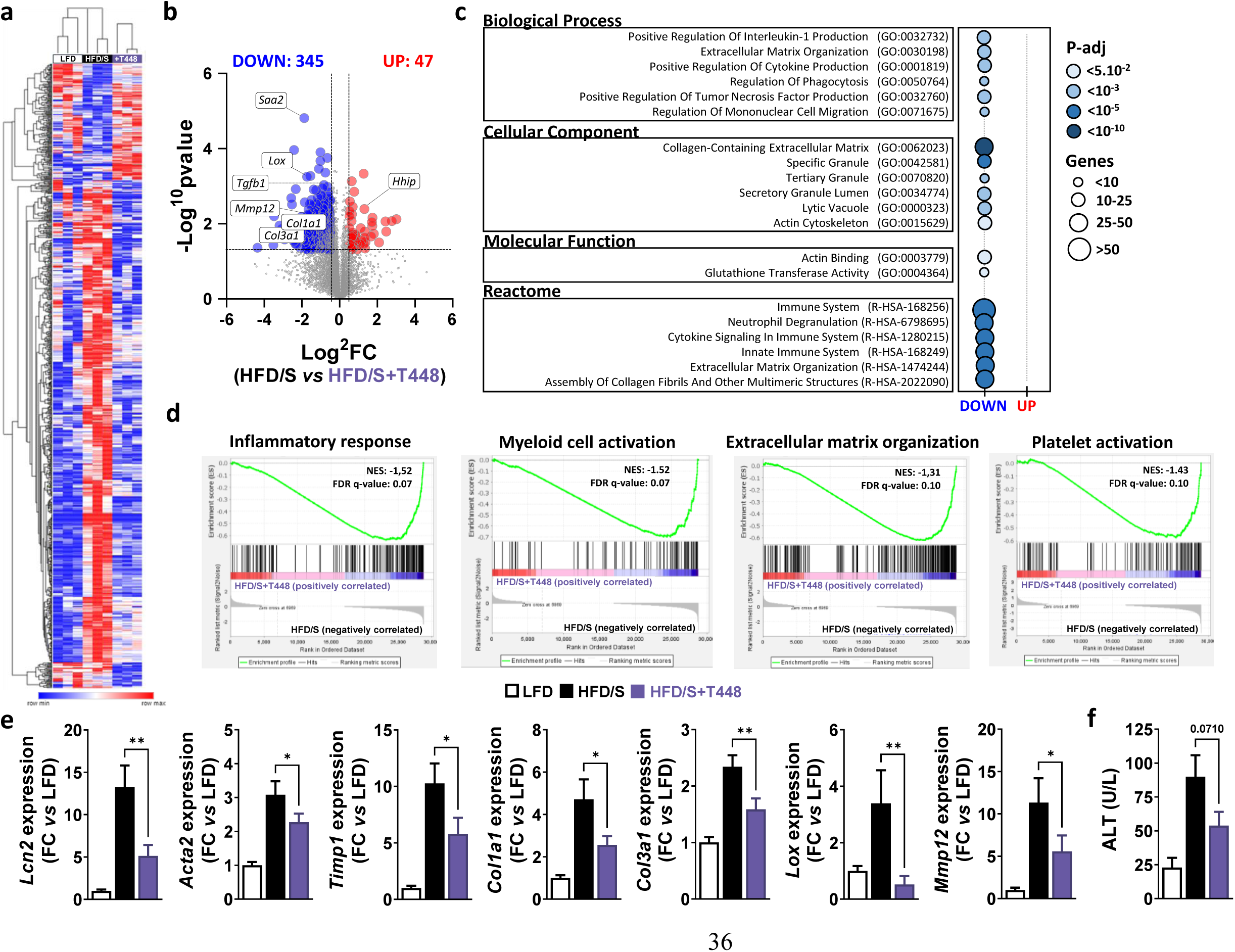
Totum-448 reduces the hepatic expression of inflammatory and fibrotic genes. LFD- and HFD/S-fed mice were treated as described in Fig. 1. Bulk RNA sequencing was performed in liver samples to assess hepatic transcriptional changes in response to T448 supplementation. Hierarchically clustered heatmap displays differentially-expressed genes (DEGs) in each group (**a**). The volcano plot depicts significantly up- and down-regulated genes in livers from HFD/S+T448-compared to HFD/S-fed mice (**b**). A GO-term analysis on DEGs (**c**) and a gene set enrichment analysis (GSEA, **d**) were performed on the whole transcriptome. Targeted qPCR was performed to assess the expression of pan-inflammatory gene *Lcn2* and fibrosis-related genes (**e**). Plasma alanine transaminase (ALT) levels were determined at week 4 in 6-hour fasted mice (**f**). Results are expressed as mean ± SEM. * p ≤ 0.05, ** p ≤ 0.01. n=3 mice per group for bulk RNA-seq (**a-d**) and n=10-12 mice per group from 2 independent experiments for targeted qPCR (**e-g**).

### Totum-448 prevents loss of tissue-resident Kupffer cells and reduces both hepatic monocyte infiltration and accumulation of pro-inflammatory monocyte-derived macrophages

To further investigate the inhibitory effect of Totum-448 on hepatic inflammation, we performed an in-depth immunophenotyping of liver leukocytes by spectral flow cytometry (see **Fig. S4a** for gating strategy). The total number of CD45^+^ hepatic leukocytes tended to be higher in HFD/S-fed mice when compared to LFD mice but was not affected by Totum-448 (**Fig. 5a**). Using Uniform Manifold Approximation and Projection for Dimension Reduction (UMAP) to visualize global changes in the major hepatic immune cell subsets (**Fig. 5b**), we observed that while Totum-448 treatment had no impact on neutrophils, NK cells, dendritic cells, and T and B cells subsets (**Fig. S4b**), it led to significant reduction of both eosinophil (**Fig. 5c**) and Ly6C^hi^ monocytes (**Fig. 5d**) in the liver from MASLD mice. Remarkably, although the total macrophage abundance was not affected in any of the groups (**Fig. 5e**), the proportion of CD11c^+^ and TREM2^+^ expressing macrophages were increased in HFD/S-fed mice and significantly lowered by Totum-448 (**Fig. 5f**), indicating a reduction in pro-inflammatory and lipid-associated macrophages, respectively. The total macrophage pool was further divided into monocyte derived CD11b^+^CLEC2^-^ macrophages (moMACS) and CD11b^low^CLEC2^+^ Kupffer cells (KCs), the latter being further divided into CLEC2^+^TIM4^-^ monocyte-derived Kupffer cells (moKCs) and resident CLEC2^+^TIM4^+^ Kupffer cells (resKCs) (**Fig. S5a**). As expected, HFD/S induced a potent loss of resKCs and a concomitant increase in moMACS in order to repopulate the KCs niche when compared to LFD-fed mice (**Fig. 5g-h**). Remarkably, Totum-448 supplementation significantly decreased both KC loss and increased accumulation of moMACS in the livers from MASLD mice (**Fig. 5g-h**), strongly suggesting a reduction in HFD/S-induced KC activation and death. Of note, an immunophenotyping was also performed in eWAT from a subset of the mice (**Fig. S5**). Totum-448, while not significantly affecting tissue leukocyte content and relative abundances of eosinophils, monocyte and T cells (**Fig. S5b-f**), may also dampen tissue inflammation by reducing tissue accumulation of both total adipose tissue macrophages (ATMs), obesity-associated pro-inflammatory CD11c^+^ATMs (**Fig. S5g-h**) and neutrophils (**Fig. S5i**).

**Figure 5.**
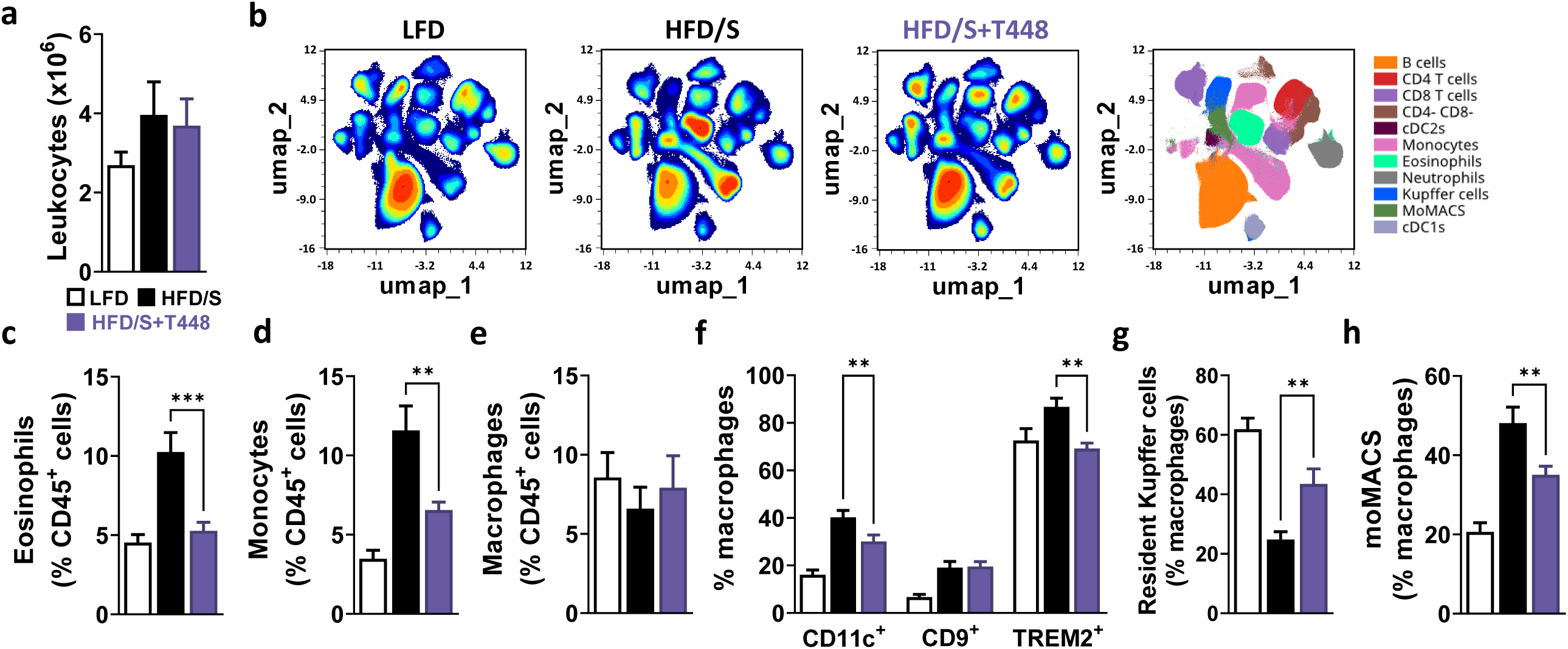
Totum-448 prevents resident Kupffer cell loss and reduces both monocyte infiltration and accumulation of pro-inflammatory monocyte-derived macrophages. LFD- and HFD/S-fed mice were treated as described in the legends of Fig. 1. The total number of CD45^+^ hepatic leukocytes was determined after isolation (**a**). Uniform Manifold Approximation and Projection for Dimension Reduction (UMAP) was used to assess global changes in the major hepatic immune cell subsets (**b**). The proportion of eosinophils (**c**) monocytes (**d**) and total hepatic macrophages (**e**) expressed as frequency of total CD45^+^ leukocytes and the proportion of CD11c^+^ and TREM2^+^ expressing macrophages were determined (**f**). The abundance of resident Kupffer cells (**g**) and monocyte-derived macrophages (moMACS, **h**) expressed as frequency of the total hepatic macrophage pool was determined. Results are expressed as mean ± SEM. * p ≤ 0.05, ** p ≤ 0.01. n=10-12 mice per group from 2 independent experiments.

## Discussion

In this study, we report and dissect the beneficial effects of Totum-448, a polyphenol-rich plant extract, on hepatic steatosis, liver inflammation and whole-body metabolic homeostasis in a dietary mouse model of MASLD.

Previous studies have shown the potential of nutraceuticals in improving cardiometabolic health, particularly in the context of insulin resistance, type 2 diabetes and MASLD^26–28,36^. Various plant-derived bioactive compounds, such as polyphenols, flavonoids, and specific fiber blends, have been shown to improve insulin sensitivity and glucose/lipid homeostasis through mechanisms independent of weight loss, notably through modulation of gut microbiota^28^.

Our results indicate that Totum-448 has a limited impact on microbiota composition in MASLD mice. While HFD/S feeding significantly altered fecal microbial diversity and composition, the marginal differences only observed in some *Firmicutes* taxa in response to Totum-448 supplementation suggest a negligible impact rather than a broad restructuring of the gut microbiome. While numerous animal studies have reported major effect of polyphenols on microbiota in a context of MASLD^37^, the relatively marginal impact of Totum-448 on microbiome composition observed in this work may be partly related to the study design. Indeed, the majority of the preclinical studies were actually carried out with the administration of polyphenols starting simultaneously with the initiation of the dietary regimen, *i.e.* assessing the impact on disease progression rather than on its regression^38^. A few studies did report significant changes in microbiota in a context of pre-established dysbiosis, but the duration of supplementation was significantly longer compared to this study^39,40^, suggesting that 4-week supplementation with Totum-448 might have been insufficient to counteract the deep-seated microbial changes induced by HFD/S.

Despite this, significant effects on hepatic steatosis were observed in Totum-448-supplemented mice. While this study was not designed to determine which specific compound was responsible for these benefits, previous research has demonstrated direct actions of certain isolated polyphenols found in Totum-448 on the liver, leading to reduced lipid accumulation. Among the most extensively studied polyphenols in HFD-fed mice, oleuropein has been shown to inhibit Wnt10b- and FGFR1-mediated signaling pathways involved in hepatic lipogenesis, while also suppressing TLR2- and TLR4-mediated pro-inflammatory signaling implicated in hepatic steatosis^41^. Additionally, it was shown to regulate lipid oxidation, lipogenesis, and inflammation via PPAR-α^42^ and activates autophagy pathways through AMPK^43^. Moreover, chlorogenic acid alleviated steatosis by inhibiting ALKBH5 activity, which in turn suppressed the ERK signaling pathway and regulated autophagy^44^. Similarly, luteolin has been reported to enhance mitochondrial biogenesis via the AMPK/PGC1α pathway, promoting fatty acid oxidation^45^. It also inhibits IL-1 and IL-18 pro-inflammatory pathways^46^ and reduces lipid accumulation by preventing LXR-mediated sterol regulatory element-binding protein-1 (SREBP-1c) activation^47^. Finally, caffeic acid has demonstrated promising hepatoprotective effects both *in vivo* and *in vitro*, reducing hepatocyte lipid accumulation and increasing autophagy, possibly through modulation of either fibroblast growth factor 21 (FGF21), FGF receptor 1 (FGFR1), β-Klotho (KLB), and/or the AMPK-SREBP-1c axis^48–50^. In addition to polyphenols, choline, another component of Totum-448, has been shown to reduce hepatic steatosis by enhancing mitochondrial function and β-oxidation while decreasing lipid accumulation^51–54^. Overall, polyphenols and choline are believed to exert anti-steatotic effects in the liver through a combination of anti-inflammatory and antioxidant mechanisms, which collectively could help alleviating insulin resistance, along with the activation of PPAR-α-mediated fatty acid oxidation and the inhibition of lipogenesis via the AMPK/SREBP-1c pathway^55,56^. In the present work, however, our liver transcriptomic analysis did not fully reflect all these pathways in the metabolic signature. Instead, the most pronounced effects were related to anti-inflammatory and anti-fibrotic responses, suggesting the participation of extrahepatic mechanisms. For instance, increased adipose tissue lipolysis due to insulin resistance is a well-recognized contributor to excessive free fatty acid delivery to the liver, resulting in steatosis. Although we did not specifically assess insulin resistance or inflammation in adipose tissue, the fact that WAT weight tended to be higher in Totum-448-supplemented mice raises intriguing possibilities for future investigations into its role in the observed metabolic effects.

In addition to its impact on metabolic homeostasis, this study provides an overview of the effects of Totum-448 on hepatic immune cell composition in HFD/S-fed mice, supporting the growing evidence that nutraceuticals, especially those of polyphenol-rich nature, can exert immunomodulatory effects which may contribute to improved metabolic outcomes^57^. In the liver, couple of key features were also associated with Totum-448 supplementation, namely reductions in HFD/S-induced eosinophilia, KC loss and moMACS accumulation. Eosinophilic inflammation has been linked to progressive MASLD and suggested to be a potential contributor to fibrotic remodeling observed in later-stage MASH^58,59^. Interestingly, we found that Totum-448 supplementation almost completely reversed HFD/S induced eosinophil accumulation indicating a potential protective effect against progressive MASH. Remarkably, Totum-448 supplementation also decreased the loss of embryonically-derived resKCs and reduced the recruitment and/or differentiation of moMACS, highlighting a beneficial remodeling of the hepatic macrophage compartment associated with reduced inflammation and MASLD progression. In addition to changes in hepatic macrophage ontogeny, modulation of their activation states have also been linked to MASLD/MASH progression, with CD11c expression being a hallmark of pro-inflammatory macrophage activation^60,61^. In our study the expression of CD11c among the total macrophage pool was significantly reduced in response to Totum-448 supplementation, indicating an overall decrease in the inflammatory activation of the hepatic macrophage compartment. Furthermore, one of the key macrophage subsets recently identified during MASLD development, called lipid-associated macrophages (LAM), were shown to arise in both WAT and the liver during obesity and to display a unique ability to store and oxidize lipids when compared to resKCs^62^. LAMs are intimately linked with fibrotic areas in the liver during MASH development and have been shown to play an essential role in regression of fibrosis^12,63^. One of the key defining markers of LAMs is the expression of Triggering receptor expressed on myeloid cells-2 (TREM2) which is strongly associated with steatohepatitis in different diet-induced murine models of MASH^59,60^. Although the exact role of TREM2^+^ macrophages in the MASLD/MASH pathophysiology still remains to be clarified, they are intimately linked with disease progression of MASLD to MASH and the rise of fibrosis and serve as an indicator of disease severity. It is however tempting to speculate that the observed decrease in TREM2^+^ macrophages induced by Totum-488 supplementation may result from a reduction in their hepatic recruitment secondary to dampening of pro-inflammatory and pro-fibrotic signaling.

In addition to the potential anti-fibrotic and immunomodulatory effects of Totum-448, the RNA sequencing data highlighted a potential lowering of platelet activation in response to Totum-448 supplementation. MASLD/MASH have been associated with a pro-thrombotic state and increased intrahepatic platelet accumulation and activation has previously been associated with various stages in MASLD and MASH pathophysiology^64–66^. For example, one study reported that platelet-derived growth factor B (PDGF-B) could activate hepatic stellate cells (HSCs) and promote liver fibrosis^64^, whilst other studies demonstrated the anti-steatotic/fibrotic effects of aspirin use^65^ and various anti-platelet drugs^67^. Platelets also directly interact with KCs, a feature that has been demonstrated in early steatosis and shown to contributes to MASH development through increased immune cell recruitment^68^. One may therefore speculate that part of the beneficial effects of Totum-448 could be related to a direct effect on intrahepatic platelet dynamics and activation, an interesting aspect to investigate that would require further studies.

Several limitations of this study ought to be acknowledged, one of them being the lack of a clear underlying mechanism explaining the observed immunometabolic effects of Totum-448. Given its polyphenol-rich composition, its benefits are likely mediated through the pleiotropic action of various bioactive molecules, influencing multiple cell types and organs both directly and indirectly. However, tissue-specific changes in insulin sensitivity were not assessed in our study, which limits our understanding of whether peripheral organs, such as the liver, adipose tissue or skeletal muscle, were specifically affected by Totum-448 supplementation. Furthermore, it is worth mentioning that the dietary model used in the current study induces a rather mild form of MASLD/MASH, with hepatic steatosis and some degree of inflammation, but without detectable fibrosis as assessed by collagen accumulation using Sirius red staining or hydroxyproline assay (data not shown). Similarly, the majority of studies describing immunological changes in the liver during MASH also rely on the use of more advanced models of MASH. One might therefore speculate that Totum-448 may eventually exert even more beneficial effects in advanced MASH stages, or could reveal stronger immunomodulatory and anti-fibrotic properties than the ones observed with the current experimental settings. Another limitation is that the study was conducted exclusively in male mice. It is well established that metabolic responses to dietary interventions, including to HFD exposure^69^ or polyphenol supplementation^70^, are exhibiting sexual dimorphism, with female mice displaying different adaptations due to hormonal variations, gut microbiota composition, immune cell profiles and intrinsic metabolic flexibility. Hence, the absence of female subjects prevents a comprehensive evaluation of whether the observed effects would be similar in both sexes. Finally, while rodent models provide valuable insights into metabolic disorders such as MASLD, their relevance to human physiology remains a key consideration. Therefore, further clinical trials are necessary to determine whether these findings translate to human populations.

In summary, we show that Totum-448 supplementation reduces both hepatic steatosis and liver inflammation, and improves whole-body metabolic homeostasis in a diet-induced MASLD mouse model. Although the underlying mechanism(s) of Totum-448 remain to be elucidated, its beneficial immunometabolic properties likely result from pleiotropic actions on various cell types and/or organs driven by a variety of plant-derived polyphenolic molecules. Altogether, supplementation with Totum-448 may constitute a promising novel nutritional approach for MASLD patients.

## Author contributions

VC, YFO, PS and BG conceptualized research; JL, MV, HJPvdZ, FO, RS and FLJ performed research; JL, MV, FO, RS, FLJ and BG analysed data; TM, MG, SLP, AZ, PS and BG supervised the study; JL, VC and BG wrote the manuscript. All authors read and approved the final manuscript.

## Acknowledgments

This study was partly funded by Valbiotis (to B.G.). The funder played no role in study design, data collection, analysis and interpretation of data, or the writing and decision to submit this manuscript.

## Data availability

The original datasets generated during the current study are available from the corresponding author on reasonable request. Raw 16S rRNA sequences are available in the NCBI data base under the bioproject number PRJNA1071341, with SRA numbers SRR27797200 to SRR27797215 and biosamples number SAMN39684451 to SAMN39684466.

