## Supplementary Data for "Totum-448, a polyphenol-rich plant extract, decreases hepatic steatosis and inflammation in diet-induced MASLD mice"

### Supplementary methods

##### ***Chemical characterization of Totum-448***

The total levels of phenolic compound (in gallic acid equivalent), sugars (in glucose equivalent), and fat (in sunflower oil equivalent) were determined in Totum-448 using Folin-Ciocalteu, Dubois, and sulfo-phospho-vanillin (SPV) colorimetric assays, respectively, as previously reported[1]. A fluorometric method using fluoraldehyde o-phthaldialdehyde reagent (OPA) was used to quantify the protein content[2], and the Lee method was used to quantify insoluble dietary fibers[3]. A more precise characterization of phytochemical compounds was performed by HPLC-UV/Visible/RID-MS using 1260 LC system and 1200 LC system with a 6110 Single Quad MS-ESI detector (Agilent Technologies, Santa Clara, CA, USA) with a C18 Prodigy reversed-phase column (250 mm × 4.6 mm, 5 µm; Phenomenex, Torrance, CA, USA) and an Atlantis HILIC Silica column (150×4.6 mm, 5 µm, Waters, The Netherlands).

##### ***Fecal microbiota analyses***

Quality filtering and processing of the raw sequencing data were done as previously described[4] using cutadapt v4.0[5] with Python v3.9.9, usearch v11.0.667[6] and mothur v1.48.0[7]. Classification of quality filtered reads was performed by comparison with the ribosomal database project (RDP trainset 14)[8,9]. Abundance tables were imported and analyzed in R v4.1. Alpha and beta diversity were estimated using package vegan 2.6-2[10] and tested using Kruskal-Wallis and MANOVA (adonis2) tests respectively. Indicator taxa analysis was performed using Kruskal-Wallis test. False Discovery Rate (FDR) correction for multiple testing was used were applicable. Indicator taxa were identified by selecting those with a significant overall association with the experimental group ($p < 0.05$), and which had at least one pairwise comparison between groups that reached significance ($p < 0.05$).

***RNA sequencing***

Sample collection and preparation

*Library preparation for Transcriptome sequencing*. A total amount of 1 µg RNA per sample was used as input material for the RNA sample preparation. Sequencing libraries were generated using NEBNext UltraTM RNA Library Prep Kit for Illumina (New England Biolabs, Ipswich, MA, USA) following manufacturer's recommendations and index codes were added to attribute sequences to each sample. Briefly, mRNA was purified from total RNA using poly-T oligo-attached magnetic beads. Fragmentation was carried out using divalent cations under elevated temperature in NEBNext First Strand Synthesis Reaction Buffer. First strand cDNA was synthesized using random hexamer primer and M-MuLV Reverse Transcriptase. Second strand cDNA synthesis was subsequently performed using DNA Polymerase I and RNase H. Remaining overhangs were converted into blunt ends via exonuclease/polymerase activities. After adenylation of 3' ends of DNA fragments, NEBNext Adaptor with hairpin loop structure were ligated to prepare for hybridization. In order to select cDNA fragments of preferentially 240 bp in length, the library fragments were purified with AMPure XP system (Beckman Coulter, Brea, CA, USA). Then 3 µl USER Enzyme (New England Biolabs, Ipswich, MA, USA) was used with size-selected, adaptor-ligated cDNA at 37°C for 15 min followed by 5 min at 95°C before PCR. PCR was then performed with Phusion High-Fidelity DNA polymerase, Universal PCR primers and Index (X) Primer. At last, PCR products were purified (AMPure XP system) and library quality was assessed on the Agilent Bioanalyzer 2100 system (Agilent Technolgies, Santa Clara, CA, USA).

*Clustering and sequencing*. The clustering of the index-coded samples was performed on a cBot Cluster Generation System using TruSeq PE Cluster Kit v4-cBot-HS (Illumia, San Diego, CA, USA) according to the manufacturer's instructions. After cluster generation, the library preparations were sequenced on an Illumina platform and paired-end reads were generated.

<u>Data analysis</u>

*Quality control*. Raw sequences (raw reads) in fastq format were first processed through in-house perl scripts. In this step, clean data (clean reads) were obtained by removing low quality reads and reads containing adapter or ploy-N from raw data. At the same time, Q20, Q30, GC-content and sequence duplication level of the clean data were calculated. Clean reads were next mapped to the reference genome sequence (*Mus musculus*, NCBI GRCm39) using the HISAT2 alignment program[11]. Only reads with a perfect match or one mismatch were further analyzed and annotated based on the reference genome.

*Gene functional annotation*. Gene function was annotated based on the following databases: Nr (NCBI non-redundant protein sequences), Nt (NCBI non-redundant nucleotide sequences), Pfam (Protein family), KOG/COG (Clusters of Orthologous Groups of proteins), Swiss-Prot (A manually annotated and reviewed protein sequence database), KO (KEGG Ortholog database) and GO (Gene Ontology).

*Quantification of gene expression levels.* Gene expression levels were estimated by fragments per kilobase of transcript per million fragments mapped (FPKM). The formula is shown as follow: FPKM = cDNA Fragments / (Mapped Fragments (Millions)) ∗ Transcript Length (kb)).

*Quantification of gene expression levels.* Differential expression analysis was performed using the DESeq2 bioconductor pipeline[12]. The resulting P-values were adjusted using the Benjamini and Hochberg's approach for controlling the false discovery rate. Genes with an adjusted P-value < 0.05 were assigned as differentially expressed.

*GO and KEGG pathway enrichment analysis.* Gene Ontology (GO) enrichment analysis of the differentially expressed genes (DEGs) was implemented by the GOseq R packages based Wallenius non-central hyper-geometric distribution， which can adjust for gene length bias in DEGs[13]. KOBAS software to test the statistical enrichment of differential expression genes in KEGG pathways[14].

***Isolation of stomal vascular fraction from adipose tissue for flow cytometry***

Epidydimal WAT were collected after a 1 min post sacrifice transcardial perfusion with PBS and further digested for isolation of stromal vascular fraction (SVF), as previously reported[15]. Briefly, eWAT samples were minced and digested for 1 hour at 37˚C under agitation (60 RPM) in 2 mL HEPES-buffered Krebs solution containing 1 mg/mL Collagenase Type I from *C. histolyticum* (Sigma-Aldrich, St. Louis, MO, USA, 125 CDU/ml), 20 mg/mL Bovine Serum Albumin (Fraction V; Sigma-Aldrich, St. Louis, MO, USA) and 6 mM D-Glucose Sigma-Aldrich, St. Louis, MO, USA ). After digestion, samples were filtered (100 µM cell strainer; Corning, NY, USA) and washed with 30 mL PBS/hiFCS/EDTA, as described above. After allowing the adipocytes to settle for 10 min, the infranatant consisting of the SVF was collected and pelleted at 350 x g for 10 min at room temperature. The pellet was treated with 1 mL erythrocyte lysis buffer, washed with 10 mL PBS/hiFCS/EDTA and counted using a hemacytometer. As for the liver, $1*10^6$ cells per sample were further processed for flow cytometry.

***Flow cytometry (eWAT leukocytes)***

Similarly to liver samples, the eWAT leucocytes were pelleted, incubated with Zombie-NIR viability dye and fixed with PFA. Post-fixation, the cells were washed with PBS/hiFSC/EDTA and incubated with a cocktail of antibodies directed against CD3, CD11b, CD11c, CD19, CD45, CD64, F4/80, Ly6C, Ly6G, NK1.1, and Siglec-F in PBS/hiFCS/EDTA supplemented with Brilliant Stain Buffer Plus (BD Biosciences, Franklin Lakes, NJ, USA) and True Stain Monocyte Blocker (Biolegend, San Diego, CA, USA) for 30 min at 4°C. After washing, as described for the liver samples, the cells were resuspended in PBS/hiFSC/EDTA and acquired on a 3-laser Cytek Aurora (Cytek Biosciences, Fremont, CA, USA).

SpectroFlo v3.0 (Cytek Biosciences, Fremont, CA, USA) was used for spectral unmixing and FlowJoTM v10.8 was used to gate the flow cytometry data for all tissues. A representative gating strategy used to gate the eWAT samples can be found in **Fig. S5**.

**Table S1**

**Chemical characterization of TOTUM-448**

| **Compound types (sorted by families)** | **Extract content (g/100 g)** |
|---|---|
| Total sugars | 27.1 |
| Total lipids | 12.6 |
| Total Proteins | 0.8 |
| Insoluble dietary fiber | 2.36 |
| Choline* | 13.67 |
| Total phenolic compounds | 8.7 |
| Total anthocyanins | 0.536 |
| Monocaffeoylquinic acids | |
| Chlorogenic acid | 0.517 |
| Cryptochlorogenic acid | 0.324 |
| Neochlorogenic acid | 0.319 |
| Other monocaffeoylquinic acids | 0.115 |
| Dicaffeoylquinic acids | |
| Cynarine | 0.229 |
| 4,5-Dicaffeoylquinic acid | 0.098 |
| 3,5-Dicaffeoylquinic acid | 0.074 |
| 3,4-Dicaffeoylquinic acid | 0.056 |
| Caffeic acid | 0.008 |
| Oleuropein | 6.223 |
| Oleuropein isomers | 0.757 |
| Ligstroside | 0.131 |
| Luteolin | 0.017 |
| Luteolin-7-O-glucoside | 0.880 |
| Luteolin-7-O-glucuronide | 0.277 |
| Luteolin-4-O-glucoside | 0.083 |
| Apigenin-7-O-glucoside | 0.062 |
| Apigenin-7-O-glucuronide | 0.139 |
| Apigenin-7-O-rutinoside | 0.037 |
| Verbascoside | 0.152 |
| Terpenes and terpenoids | |
| Oleanolic acid | 0.199 |
| Cynaropicrin | 0.139 |
| Saponins | |
| Chrysanthellin A | 0.133 |
| Chrysanthellin B | 0.215 |
| Alkaloids | |
| Piperin | 0.044 |

* in choline chloride equivalent

#### Table S2

| Gene | Accession number | Forward primer | Reverse primer |
|---|---|---|---|
| *Acta2* | NM_007392.3 | AGCCATCTTTCATTGGGATGG | CCCCTGACAGGACGTTGTTA |
| *Col1a1* | NM_007742.3 | GAGAGGTGAACAAGGTCCCG | AAACCTCTCTCGCCTCTTGC |
| *Col3a1* | NM_009930 | TGACTGTCCCACGTAAGCAC | GAGGGCCATAGCTGAACTGA |
| *Lcn2* | NM_008491 | GCCACTCCATCTTTCCTGTTG | AAGAGGCTCCAGATGCTCCTT |
| *Lox* | NM_010728.3 | CACTGCACACACACAGGGAT | ATTGTGCAGCCTGAGGCATA |
| *Mmp12* | NM_008605.3 | GAACTTGCAGTCGGAGGGAA | TCTTGACAAGTACCATTCAGCA |
| *Rplp0* | NM_007475 | TCTGGAGGGTGTCCGCAACG | GCCAGGACGCGCTTGTACCC |
| *Timp1* | NM_011593 | AGTGCCTGCAGCTTCTTGGT | CAGCCAGCACTATAGGTCTTTGAG |

#### Table S3

| Specificity | Clone | Conjugate | Supplier | Catalog # | Titration |
|---|---|---|---|---|---|
| CD3 | 17A2 | APC-Fire 810 | Biolegend | 100268 | 1:100 |
| CD3 | 17A2 | BV605 | Biolegend | 100237 | 1:200 |
| CD4 | RM4-5 | BV650 | Biolegend | 100546 | 1:200 |
| CD8 | 53-6.7 | BV711 | Biolegend | 100748 | 1:200 |
| CD11b | M1/70 | PE-Cy7 | eBioscience | 25-0112-82 | 1:6000 |
| CD11c | HL3 | V450 | BD | 560521 | 1:100 |
| CD19 | MB19-1 | FITC | Biolegend | 101506 | 1:100 |
| CD19 | 1D3 | BV480 | BD | 566107 | 1:400 |
| CD45 | 30-F11 | BV785 | Biolegend | 103149 | 1:800 |
| CD64 | X54-5/7.1 | PE-Dazzle 594 | Biolegend | 139320 | 1:100 |
| CD90.2 | 30-H12 | Alexa Fluor 700 | Biolegend | 105320 | 1:1600 |
| CLEC2 | 17D9 | FITC | BioRad | MCA5700 | 1:400 |
| F4.80 | BM8 | BV711 | Biolegend | 123147 | 1:200 |
| Ly6C | HK1.4 | APC-Cy7 | Biolegend | 128026 | 1:800 |
| Ly6C | HK1.4 | BV510 | Biolegend | 128033 | 1:800 |
| Ly6G | 1A8 | BV650 | Biolegend | 127641 | 1:800 |
| NK1.1 | PK136 | PerCP-Cy5.5 | Biolegend | 108728 | 1:100 |
| Siglec-F | E50-2440 | PE | BD | 552126 | 1:200 |
| TIM4 | RMT4-54 | PerCP-eFluor710 | eBioscience | 46-5866-82 | 1:1000 |
| TREM2 | 6E9 | PE | Biolegend | 8244805 | 1:100 |
| **Other reagents** | | | | | |
| Viability | | Zombie-NIR | Biolegend | 423106 | 1:1000 |
| True Stain Monocyte blocker | | | Biolegend | 426103 | 1:20 |
| Brilliant Stain Buffer Plus | | | BD | 566385 | 1:10 |

#### Table S4

| | Mean +/- SD | | | p-values | | | |
|---|---|---|---|---|---|---|---|
| **Phylum** | **LFD** | **HFD/S** | **HFD/S+T448** | **Overall** | **LFD vs HFD/S** | **LFD vs HFD/S+T448** | **HFD/S vs HFD/S+T448** |
| **Bacteroidetes** | 18.67% +/- 1.24% | 25.39% +/- 3.47% | 29.24% +/- 2.86% | p= 0.0049 ** | p= 0.0104 * | p= 0.0105 * | p= 0.0881 ns |
| **Actinobacteria** | 23.93% +/- 5.89% | 6.71% +/- 3.75% | 5.28% +/- 6.04% | p= 0.0046 ** | p= 0.0039 ** | p= 0.0105 * | p= 0.5212 ns |
| **Tenericutes** | 0.78% +/- 0.84% | 0.02% +/- 0.05% | 0.00% +/- 0.01% | p= 0.0030 ** | p= 0.0033 ** | p= 0.0096 ** | p= 0.6926 ns |

#### Table S5

| Phylum | Class | Order | Family | Genus | Mean +/- SD | | | p-values | | |
|---|---|---|---|---|---|---|---|---|---|---|
| | | | | | LFD | HFD/S | HFD/S+T448 | Overall | LFD vs HFD/S | HFD/S vs HFD/S+T448 |
| **Firmicutes** | Erysipelotrichia | Erysipelotrichales | Erysipelotrichaceae | Allobaculum | 0.22% +/- 0.07% | 0.79% +/- 0.42% | 0.14% +/- 0.02% | p= 0.0021 ** | p= 0.0064 ** | p= 0.0103 * |
| **Bacteroidetes** | Bacteroidia | Bacteroidales | Rikenellaceae | Alistipes | 3.42% +/- 2.21% | 7.43% +/- 2.82% | 7.13% +/- 1.81% | p= 0.0337 * | p= 0.0250 * | p= 1.0000 ns |
| **Firmicutes** | Clostridia | Clostridiales | Lachnospiraceae | Clostridium_XlVa | 0.56% +/- 0.25% | 1.19% +/- 0.36% | 1.93% +/- 0.70% | p= 0.0119 * | p= 0.0247 * | p= 0.0871 ns |
| **Actinobacteria** | Actinobacteria | Bifidobacteriales | Bifidobacteriaceae | uncl. Bifidobacteriaceae | 1.33% +/- 0.22% | 0.68% +/- 0.36% | 0.50% +/- 0.58% | p= 0.0131 * | p= 0.0104 * | p= 0.6698 ns |
| **Firmicutes** | Erysipelotrichia | Erysipelotrichales | Erysipelotrichaceae | Clostridium_XVIII | 0.01% +/- 0.01% | 0.09% +/- 0.09% | 0.06% +/- 0.05% | p= 0.0227 * | p= 0.0275 * | p= 0.4542 ns |
| **Actinobacteria** | Actinobacteria | Bifidobacteriales | Bifidobacteriaceae | Bifidobacterium | 20.45% +/- 4.44% | 4.61% +/- 3.24% | 3.87% +/- 5.57% | p= 0.0046 ** | p= 0.0039 ** | p= 0.5224 ns |
| **Bacteroidetes** | Bacteroidia | Bacteroidales | uncl. Bacteroidales | uncl. Bacteroidales | 0.66% +/- 0.12% | 4.66% +/- 2.71% | 1.69% +/- 0.38% | p= 0.0040 ** | p= 0.0039 ** | p= 0.2864 ns |
| **Bacteroidetes** | Bacteroidia | Bacteroidales | Muribaculaceae | Muribaculum | 0.80% +/- 0.19% | 0.46% +/- 0.06% | 0.45% +/- 0.13% | p= 0.0159 * | p= 0.0080 ** | p= 0.8307 ns |
| **Firmicutes** | Bacilli | Lactobacillales | Streptococcaceae | Lactococcus | 0.00% +/- 0.01% | 0.71% +/- 0.63% | 1.59% +/- 0.21% | p= 0.0023 ** | p= 0.0028 ** | p= 0.0881 ns |
| **Tenericutes** | Mollicutes | Anaeroplasmatales | Anaeroplasmataceae | Anaeroplasma | 0.78% +/- 0.84% | 0.02% +/- 0.05% | 0.00% +/- 0.01% | p= 0.0030 ** | p= 0.0033 ** | p= 0.6926 ns |
| **Firmicutes** | uncl. Firmicutes | uncl. Firmicutes | uncl. Firmicutes | uncl. Firmicutes | 1.17% +/- 0.26% | 1.57% +/- 0.27% | 0.83% +/- 0.36% | p= 0.0172 * | p= 0.0370 * | p= 0.0176 * |
| **Firmicutes** | Bacilli | Lactobacillales | Lactobacillaceae | Lactobacillus | 0.34% +/- 0.36% | 1.70% +/- 0.98% | 0.17% +/- 0.17% | p= 0.0209 * | p= 0.0250 * | p= 0.0190 * |
| **Firmicutes** | Bacilli | Lactobacillales | Lactobacillaceae | Limosilactobacillus | 0.13% +/- 0.27% | 0.63% +/- 0.37% | 0.08% +/- 0.12% | p= 0.0170 * | p= 0.0156 * | p= 0.0187 * |
| **Firmicutes** | Bacilli | Lactobacillales | Lactobacillaceae | uncl. Lactobacillaceae | 0.14% +/- 0.12% | 0.42% +/- 0.20% | 0.12% +/- 0.15% | p= 0.0299 * | p= 0.0198 * | p= 0.0543 ns |
| **Firmicutes** | Bacilli | uncl. Bacilli | uncl. Bacilli | uncl. Bacilli | 0.06% +/- 0.05% | 0.22% +/- 0.12% | 0.06% +/- 0.03% | p= 0.0493 * | p= 0.0303 * | p= 0.0521 ns |
| **Bacteroidetes** | Bacteroidia | Bacteroidales | Muribaculaceae | uncl. Muribaculaceae | 7.86% +/- 0.88% | 5.41% +/- 1.64% | 7.07% +/- 1.30% | p= 0.0141 * | p= 0.0065 ** | p= 0.0881 ns |
| **Firmicutes** | Bacilli | Lactobacillales | uncl. Lactobacillales | uncl. Lactobacillales | 0.07% +/- 0.03% | 0.18% +/- 0.10% | 0.09% +/- 0.04% | p= 0.0175 * | p= 0.0071 ** | p= 0.0513 ns |
| **Firmicutes** | Erysipelotrichia | Erysipelotrichales | Erysipelotrichaceae | uncl. Erysipelotrichaceae | 29.63% +/- 3.91% | 29.26% +/- 6.36% | 14.10% +/- 4.33% | p= 0.0129 * | p= 0.5218 ns | p= 0.0105 * |
| **Bacteroidetes** | Bacteroidia | Bacteroidales | Porphyromonadaceae | Parabacteroides | 0.37% +/- 0.20% | 0.61% +/- 0.56% | 2.07% +/- 1.07% | p= 0.0269 * | p= 0.8717 ns | p= 0.0330 * |
| **Actinobacteria** | Coriobacteriia | Eggerthellales | Eggerthellaceae | uncl. Eggerthellaceae | 0.00% +/- 0.01% | 0.00% +/- 0.00% | 0.07% +/- 0.06% | p= 0.0179 * | p= 0.3173 ns | p= 0.0182 * |
| **Firmicutes** | Clostridia | Clostridiales | Ruminococcaceae | Neglecta | 0.07% +/- 0.03% | 0.04% +/- 0.03% | 0.15% +/- 0.06% | p= 0.0081 ** | p= 0.1000 ns | p= 0.0098 ** |
| **Firmicutes** | Clostridia | Clostridiales | Lachnospiraceae | Schaedlerella | 0.00% +/- 0.00% | 0.10% +/- 0.17% | 0.40% +/- 0.34% | p= 0.0164 * | p= 0.1530 ns | p= 0.0842 ns |

#### Supplementary figure legends

**Figure S1. Totum-448 induces a dose-dependent decrease in bodyweight.** 10 week-old C57BL/6JOlaHsd male mice were fed either a low-fat diet (LFD, open squares/bars) or high-fat diet (HFD) supplemented with sucrose in the drinking water (10% w/v, HFD/S) for a period of 12 weeks after which the HFD was either supplemented with Totum-448 (T448, 1.5% (circles), 2% (diamonds) or 2.5% (triangles) g/g; purple symbols/bars) or left without supplementation (control; black squares/bars) for 4 additional weeks (**a**). Bodyweight was monitored throughout the supplementation period and the total body weight loss was determined at week 4 (**b-c**). The cumulative food intake and mean daily food intake were determined (**d**). The cumulative liquid energy intake and mean daily liquid energy intake were calculated (**e**). Combined energy intake from both food and liquid consumption were calculated and presented as cumulative energy intake and mean daily energy intake (**f**). Mean daily feces production was calculated (**g**). Results are expressed as mean ± SEM. * $p \leq 0.05$, **** $p \leq 0.0001$. n=5-6 mice per group (**a-c**); n=2-3 cages per group (**d-g**).

**Figure S2. T448 already improves whole-body insulin sensitivity after 2 weeks of supplementation.** At week 2 of treatment, body weight (**a**) and body composition (**b**) were determined. The fasting glucose and insulin levels (**c-d**) were determined and used to calculate HOMA-IR (**e**). Results are expressed as mean ± SEM. ** $p \leq 0.01$. n=10-12 mice per group from 2 independent experiments.

**Figure S3. Totum-448 does not have significant impact on blood leukocytes.** LFD- and HFD/S-fed mice were treated as described in Fig. 1a. Blood was collected at sacrifice and leukocyte composition was analyzed by spectral flow cytometry. A representative gating strategy used to identify the different blood leukocyte subsets is depicted (**a**). The number of leukocytes per ml of blood was calculated (**b**). Uniform Manifold Approximation and Projection for Dimension Reduction (UMAP) was used to globally assess the composition of the circulating immune cell compartment (**c**). The total number of circulating B cells, NK cells, $CD4^+$ and $CD8^+$ T cells, eosinophils, neutrophils and monocytes were calculated (**d**). Results are expressed as mean ± SEM. * $p \leq 0.05$. n=10-12 mice per group from 2 independent experiments.

**Figure S4. Effect of Totum-448 on hepatic immune cell subsets.** LFD- and HFD/S-fed mice were treated as described in Fig. 1a. A representative gating strategy used to identify various hepatic immune cell subsets is shown (**a**). The relative proportion of neutrophils, NK cells, dendritic cells (DCs), T cells and B cells were determined and expressed as percentage of the total $CD45^+$ leukocyte pool. Results are expressed as mean ± SEM. *** $p \leq 0.001$. n=10-12 mice per group from 2 independent experiments.

**Figure S5. Effect of Totum-448 on immune cell subsets in epididymal white adipose tissue.** LFD- and HFD/S-fed mice were treated as described in Fig. 1a. A representative gating strategy used to identify various immune cell subsets in epididymal white adipose tissue (eWAT) is shown (**a**). The total number of eWAT $CD45^+$ leukocytes was determined after isolation (**b**). Uniform Manifold Approximation and Projection for Dimension Reduction (UMAP) was used to globally assess the composition of the eWAT immune compartment (**c**). The relative proportion of eosinophils (**d**), monocytes (**e**), T cells (**f**), total adipose tissue macrophages (ATMs, **g**), $CD11c^+$ ATMs (**h**), neutrophils (**i**) and were determined and expressed as frequency of the total $CD45^+$ leukocyte pool or as frequency of total ATMs. Results are expressed as mean

± SEM. * $p \leq 0.05$, ** $p \leq 0.01$. n=3-4 mice per group.

**Figure S1**

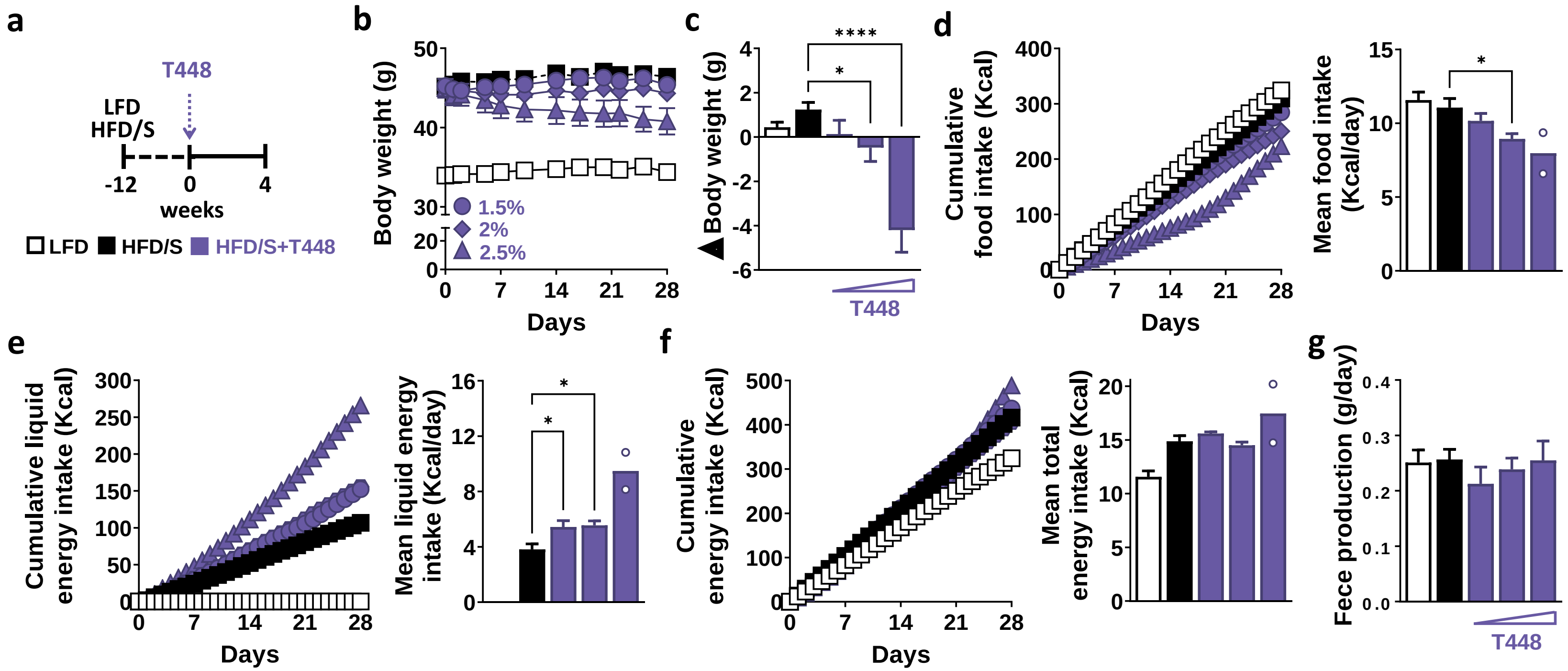
a
T448
LFD
HFD/S
-12
0
4
weeks
LFD
HFD/S
HFD/S+T448
b
Body weight (g)
1.5%
2%
2.5%
Days
c
Δ Body weight (g)
****
*
T448
d
Cumulative food intake (Kcal)
Days
Mean food intake (Kcal/day)
*
e
Cumulative liquid energy intake (Kcal)
Days
Mean liquid energy intake (Kcal/day)
*
*
f
Cumulative energy intake (Kcal)
Days
Mean total energy intake (Kcal)
g
Fece production (g/day)
T448

**Figure S2**

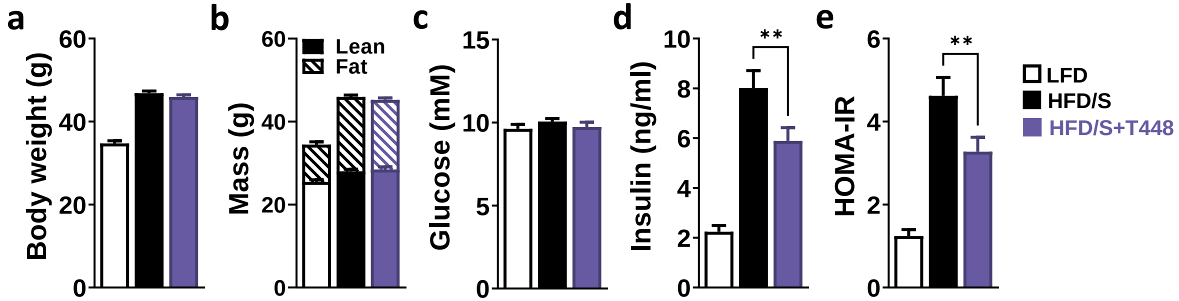
a
Body weight (g)
60
40
20
0
b
Mass (g)
Lean
Fat
60
40
20
0
c
Glucose (mM)
15
10
5
0
d
Insulin (ng/ml)
10
8
6
4
2
0
**
e
HOMA-IR
6
4
2
0
**
LFD
HFD/S
HFD/S+T448

**Figure S3**

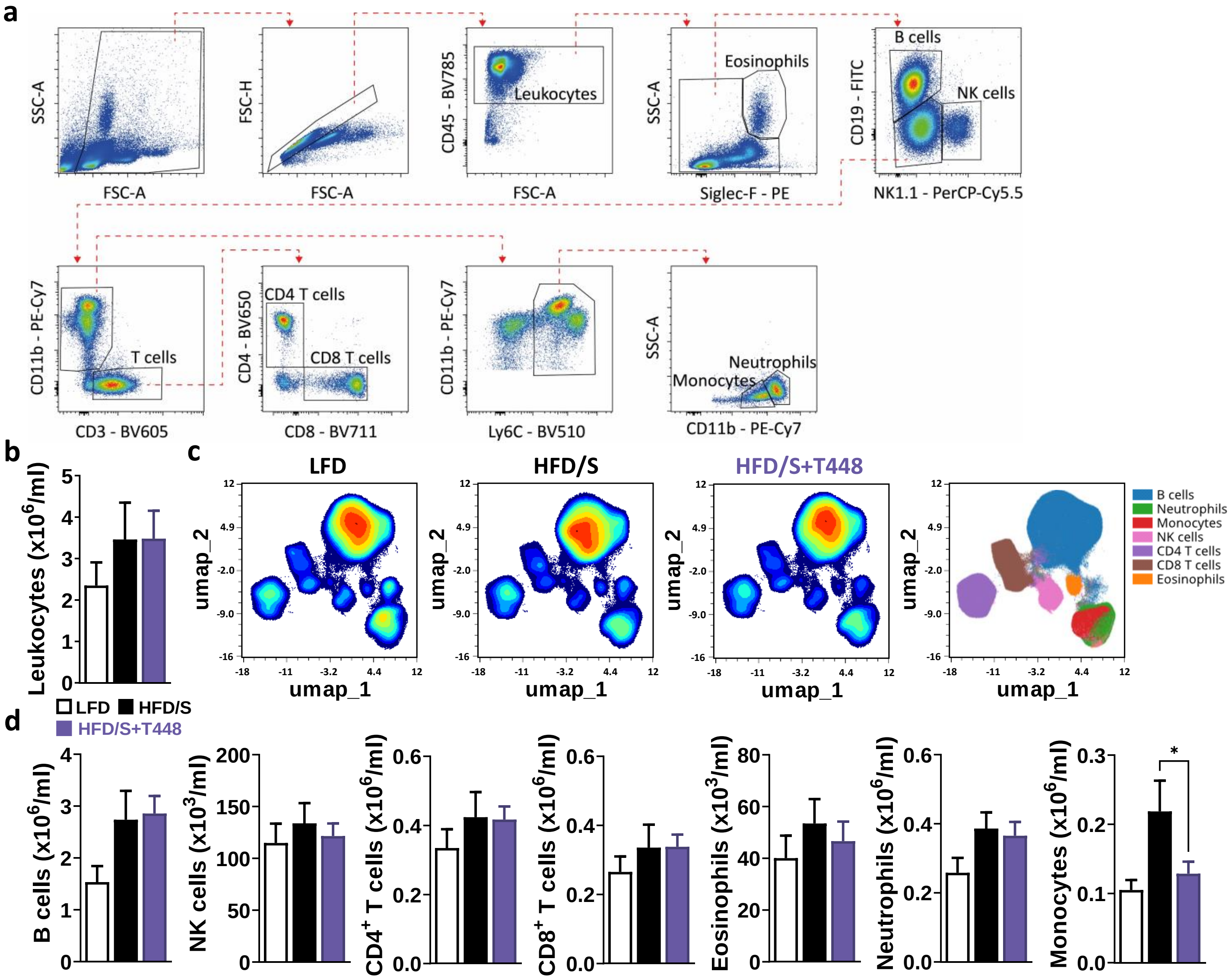
a
SSC-A
FSC-A
FSC-H
FSC-A
CD45 - BV785
Leukocytes
FSC-A
SSC-A
Eosinophils
Siglec-F - PE
CD19 - FITC
B cells
NK cells
NK1.1 - PerCP-Cy5.5
CD11b - PE-Cy7
T cells
CD3 - BV605
CD4 - BV650
CD4 T cells
CD8 T cells
CD8 - BV711
CD11b - PE-Cy7
Ly6C - BV510
SSC-A
Neutrophils
Monocytes
CD11b - PE-Cy7
b
Leukocytes (x10^6/ml)
c
LFD
HFD/S
HFD/S+T448
umap_2
umap_1
B cells
Neutrophils
Monocytes
NK cells
CD4 T cells
CD8 T cells
Eosinophils
LFD
HFD/S
HFD/S+T448
d
B cells (x10^6/ml)
NK cells (x10^3/ml)
CD4+ T cells (x10^6/ml)
CD8+ T cells (x10^6/ml)
Eosinophils (x10^3/ml)
Neutrophils (x10^6/ml)
Monocytes (x10^6/ml)
*

**Figure S4**

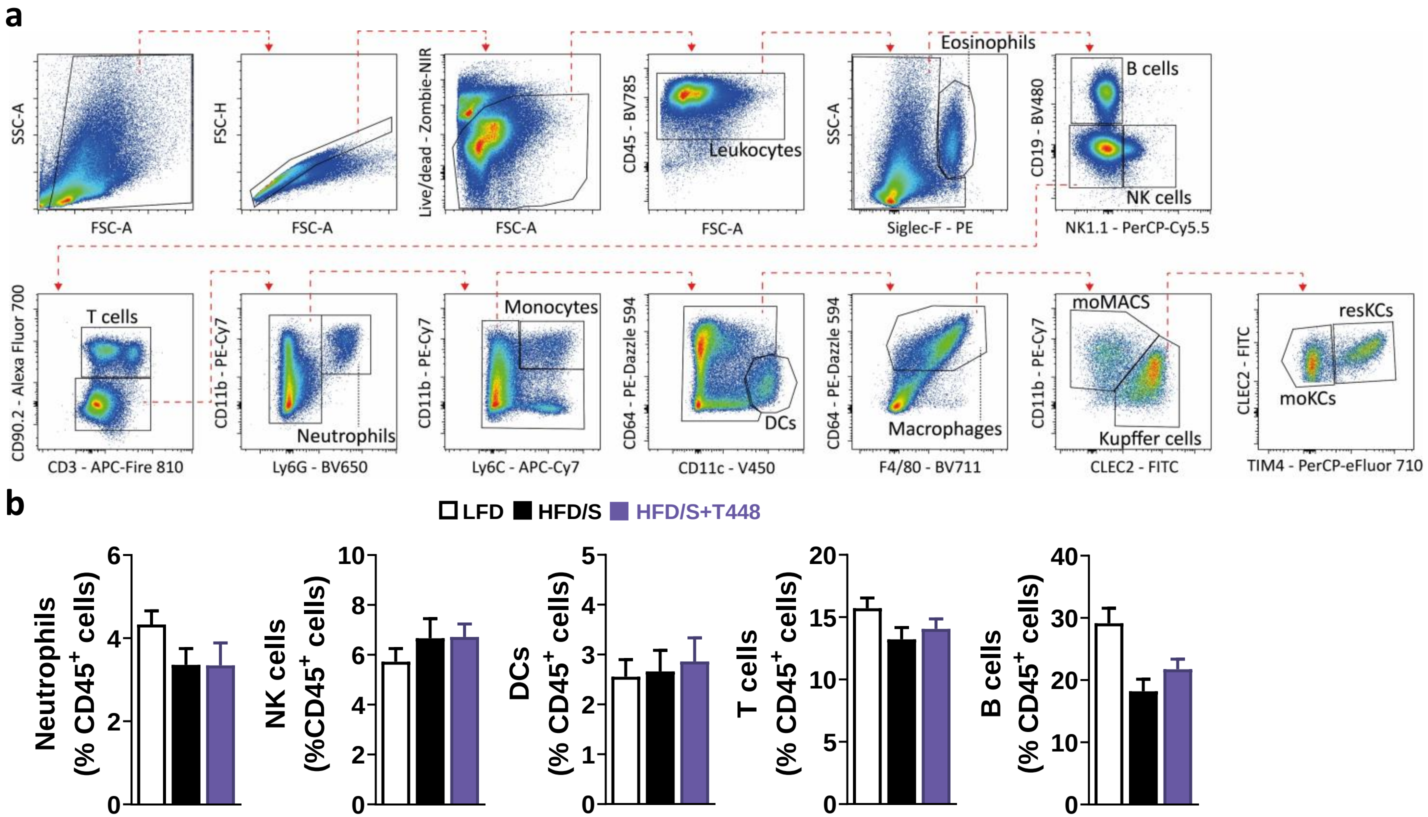
a
SSC-A
FSC-A
FSC-H
FSC-A
Live/dead - Zombie-NIR
FSC-A
CD45 - BV785
Leukocytes
FSC-A
SSC-A
Eosinophils
Siglec-F - PE
CD19 - BV480
B cells
NK cells
NK1.1 - PerCP-Cy5.5
CD90.2 - Alexa Fluor 700
T cells
CD3 - APC-Fire 810
CD11b - PE-Cy7
Neutrophils
Ly6G - BV650
CD11b - PE-Cy7
Monocytes
Ly6C - APC-Cy7
CD64 - PE-Dazzle 594
DCs
CD11c - V450
CD64 - PE-Dazzle 594
Macrophages
F4/80 - BV711
CD11b - PE-Cy7
moMACS
Kupffer cells
CLEC2 - FITC
CLEC2 - FITC
resKCs
moKCs
TIM4 - PerCP-eFluor 710
b
LFD
HFD/S
HFD/S+T448
Neutrophils (% CD45+ cells)
NK cells (%CD45+ cells)
DCs (% CD45+ cells)
T cells (% CD45+ cells)
B cells (% CD45+ cells)

#### Figure S5

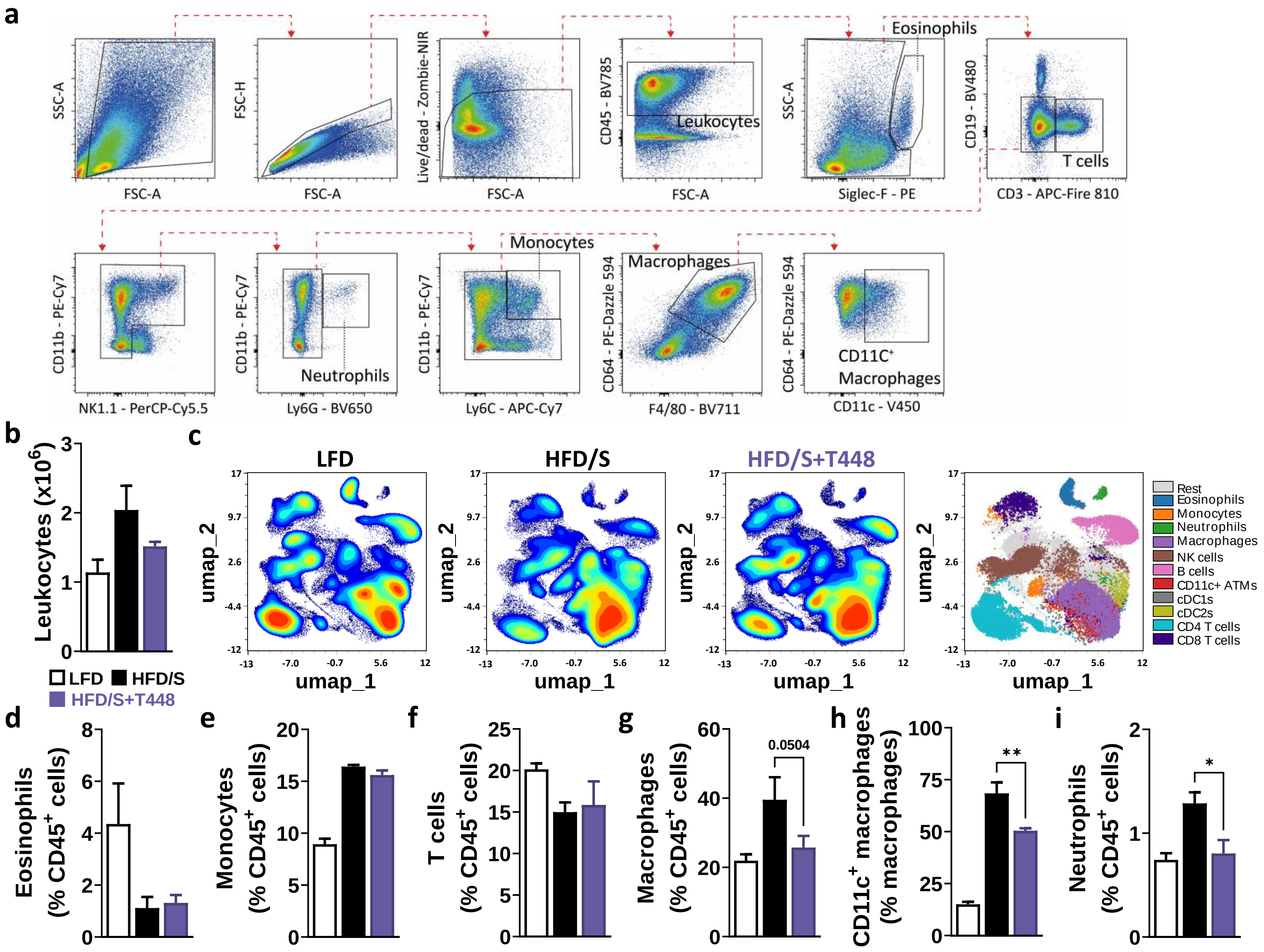
a
SSC-A
FSC-A
FSC-H
FSC-A
Live/dead - Zombie-NIR
FSC-A
CD45 - BV785
Leukocytes
FSC-A
SSC-A
Eosinophils
Siglec-F - PE
CD19 - BV480
T cells
CD3 - APC-Fire 810
CD11b - PE-Cy7
NK1.1 - PerCP-Cy5.5
CD11b - PE-Cy7
Neutrophils
Ly6G - BV650
CD11b - PE-Cy7
Monocytes
Ly6C - APC-Cy7
CD64 - PE-Dazzle 594
Macrophages
F4/80 - BV711
CD64 - PE-Dazzle 594
CD11C+ Macrophages
CD11c - V450
b
Leukocytes (x10^6)
LFD
HFD/S
HFD/S+T448
c
LFD
HFD/S
HFD/S+T448
umap_2
umap_1
Rest
Eosinophils
Monocytes
Neutrophils
Macrophages
NK cells
B cells
CD11c+ ATMs
cDC1s
cDC2s
CD4 T cells
CD8 T cells
d
Eosinophils (% CD45+ cells)
e
Monocytes (% CD45+ cells)
f
T cells (% CD45+ cells)
g
Macrophages (% CD45+ cells)
0.0504
h
CD11c+ macrophages (% macrophages)
**
i
Neutrophils (% CD45+ cells)
*
